# Bcl-2 Oligomerizes Bax on the Mitochondrial Membrane Surface Preventing the Initial Stages of Apoptosis

**DOI:** 10.1101/2025.01.24.634681

**Authors:** Sophie E. Ayscough, Luke A. Clifton, Jörgen Ådén, Sebastian Köhler, Nicolò Paracini, James Doutch, Éilís C Bragginton, Anna. E. Leung, Oliver Bogojevic, Jia-Fei Poon, Tamás Milán Nagy, Hanna P. Wacklin-Knecht, Gerhard Gröbner

## Abstract

The Bcl-2 family of proteins control mitochondrial outer membrane (MOM) pore formation, crucial to cellular clearance *via* apoptosis. However, the molecular principles by which opposing family members inhibit, mediate or promote MOM perforation remain elusive. Here, we demonstrate that cell-protecting Bcl-2 directly sequesters cell-killing Bax into a protein-protein complex at the membrane interface preventing Bax forming apoptotic pores. Neutron reflectometry showed Bax association with Bcl-2 occurs through the formation of Bax protein oligomers on the membrane surface. Bax binding to the membrane surface was proportional to the membrane-embedded Bcl-2 suggestive of protein-protein complex formation through both Bcl-2/Bax and Bax/Bax interactions. Bcl-2/Bax sequestration to prevent perforation was observed in membrane models with and without pro-apoptotic cardiolipin present. Our findings shed fundamental new light on the communication of Bcl-2 with its cell-killing relatives at the mitochondriás membraneous exterior to prevent cells from undergoing apoptosis.

**Teaser:** *Membrane-embedded Bcl-2 sequesters Bax, to prevent the perforation of mitochondrial membranes by Bax which would initiate apoptosis*.

## Introduction

Apoptosis is a form of regulated cell death that is essential for human development and health (*1*). Upon activation of the intrinsic apoptotic pathway the progression towards cellular clearance *via* inducing controlled cellular death requires the intimate involvement of the celĺs powerhouse, the mitochondrion (*2, 3*). During this process the mitochondriońs protective shell, namely the mitochondrial outer membrane (MOM), undergoes permeabilization, thereby releasing apoptotic factors including cytochrome c. This finally triggers an irreversible signaling cascade, causing the death of the cell (*3, 4*). To avoid undesired clearance of healthy cells, this pathway is critically controlled by a sophisticated network of opposing members of the Bcl-2 (B-cell CLL/lymphoma-2) protein family. This family consists, besides a group of simple BH3-only domain initiators (pro-apoptotic/cell-killing), of two main groups of multi-domain Bcl-2 proteins called guardians (anti-apoptotic/pro-survival) and executioners (pro-apoptotic/cell-killing) (*2, 3*). To control any apoptotic activity these opposing family members meet at the MOM where they interact with each other to determine a cell’s fate: survival by assuring MOM integrity or cell death by inducing membrane leakage (*4, 5*). However, any failure in this communication can have severe consequences including e.g. embryonal defects, heart diseases, neurodegenerative disorders and most prominently cancer (*6–8*). In nearly 50% of all human cancers, overproduction of the cell-protecting members like the Bcl-2 protein, that gives its name to the family, promotes tumor development and therapy resistance by lowering their susceptibility to apoptosis (*6, 9*). On the contrary, misfolded aggregation-prone proteins might trigger premature neuronal death in neurodegenerative diseases by inhibiting Bcl-2 proteins; acting thereby as neurotoxic agents (*10*).

In general, clearance of undesired cells is initiated by apoptotic stress signals causing the massive recruitment of pro-apoptotic Bcl-2 members such as Bax (Bcl-2 associated protein X) to the MOM (*2, 11–14*). There, activated Bax can oligomerize and generate pores in the membrane (*12, 14–16*), enabling the release of apoptotic factors to trigger further steps of the apoptotic cascade towards irreversible cell destruction. To ensure tissue homeostasis by protecting healthy cells from undesired demise, guardians such as the Bcl-2 protein itself preserve mitochondrial integrity through sequestration of activated Bax and its relatives (*2*), a mechanism also exploited by cancer cells to ensure survival (*17, 18*). However, the molecular mechanism and principles by which the most prominent family representatives Bcl-2 and Bax are organized at the MOM and interact with each other, to control apoptotic pore formation, remains elusive due to the lack of molecular and structural insight in the membrane environment.

Despite the absence of an atomic resolution model for membrane perforating Bax and its assemblies, numerous studies provide a basic molecular understanding of the ability of Bax to recognize MOMs and create pores. These studies unraveled not only the spatial and temporal organization of Bax at the MOM during the pore formation process but also the active roles of lipids in pore formation and subsequent release of apoptotic factors (*12, 14–16*). However, for Bcl-2 protein when active in its native MOM environment, there is no reliable molecular model for how it recognizes and inhibits Bax to prevent pore formation. The main reasons for the absence of insight are the lack of atomic-level information about the structure of full-length Bcl-2 in a membrane and its structural plasticity. Bcl-2, as a typical “tail anchored membrane protein” has an amphitropic globular domain forming its extended binding groove and a transmembrane anchoring domain at its C-terminal (*19–21*). While it is thought that these features allow Bcl-2 to be initially loosely membrane anchored with its globular body exposed to the cytosol, recent work by us using neutron reflectometry and NMR demonstrated that Bcl-2 is embedded in the mitochondrial membrane (*22, 23*). This membrane embedded location of Bcl-2 is likely required for Bcl-2 to recognize and inhibit pro-apoptotic Bcl-2 family members upon their membrane association and partial penetration; this has already been predicted by Hill *et. Al* (*20*) to be necessary for Bcl-2 to exert its protective function in the MOM. Based on our recent work on Bax membrane perforation(*14, 22, 23*), our main aim here is to provide a fundamental molecular-level view into the mechanism by which Bcl-2 recognizes and sequesters Bax at the mitochondrial outer membrane to protect cells.

By using our recently established complementary neutron reflectometry (NR) and attenuated total reflection Fourier-Transform infrared spectroscopy (ATR-FTIR) approach (*14*), we followed the spatial and temporal fate of the Bcl-2 and Bax proteins during their interplay at their target membrane environment. NR in combination with isotope contrast labelling of individual components provides an in-depth molecular insight across membranes providing locations and relative distributions of the lipids and proteins (*24, 25*). The method is ideal for following the membrane interactions of Bax in the absence and presence of Bcl-2. By *in-situ* titration of Bax to model MOM bilayers either containing or lacking Bcl-2 we were able to monitor Bax-membrane and Bax/Bcl-2 interactions, and any potential membrane damage as caused by Bax alone (*14*).Time dependent changes in the location and distribution of the various protein and lipid components in the membrane system can be determined from the NR-derived scattering length density profiles (*22*) and the associated component volume fraction profiles (*26*). In addition, ATR-FTIR enabled us to extract, on a faster time scale, the membrane-association kinetics of Bax, and the behavior of the proteins and matrix forming lipids by detecting protein and lipid specific infrared spectroscopy signals (*14, 25*). Finally, electron microscopy (EM) imaging of Bcl-2 containing vesicles was used to complement NR data showing the changes in membrane morphology before and after the interaction of Bax.

## Results and Discussion

Neutron reflectometry (NR) was used to provide membrane level insight into the molecular details by which Bcl-2 inhibits Bax to prevent apoptotic pore formation. Vesicle fusion was used to deposit a series of MOM models on silicon substrate surfaces. The interactions of Bax with these MOM models with and without the pro-survival Bcl-2 were then structurally compared. The MOM models were supported lipid bilayers (SLBs) composed of either POPC (1-palmitoyl-2-oleoyl-glycero-3-phosphocholine) only or POPC and 10% (mol/mol) cardiolipin (CL), with varying amounts of embedded Bcl-2. The different models were then used to examine how the presence of Bcl-2 changes the interaction of Bax with the MOM when both a pro-apoptotic lipid (CL) (*27*) and anti-apoptotic Bcl-2 (*28*) were simultaneously present.

### Bax forms pores in POPC only membranes

Control NR measurements on basic POPC bilayers revealed that Bax induced pore formation into the SLB through lipid removal and redistribution of this into protein-lipid clusters on the membrane surface (Fig 1). The final POPC-Bax dataset was obtained at 30°C upon equilibrium Bax interaction (i.e. when no further time dependent changes were observed in the NR data). This mechanism was similar, but significantly slower (see supporting information Fig S8), than that previously observed by us in the presence of CL containing target membranes (*14*). However, a higher volume fraction of Bax was found to embed within the hydrophobic core of the POPC lipid bilayer compared to its interaction with CL containing membranes. Additionally, our previous results showed that lipid removed from the CL containing MOM-models was completely redistributed to the membrane surface in Bax-lipid clusters. For the Bax interaction with the POPC only membrane this was only partially found with a ∼20% loss of lipids (see Table 1) from the bilayer during Bax induced pore formation but only around 25% of that removed lipid appeared in Bax lipid complexes found on the membrane surface, suggesting the other 75% of the removed lipids moved into the bulk solution possibly as solution stable Bax-POPC clusters.

**Figure 1.**
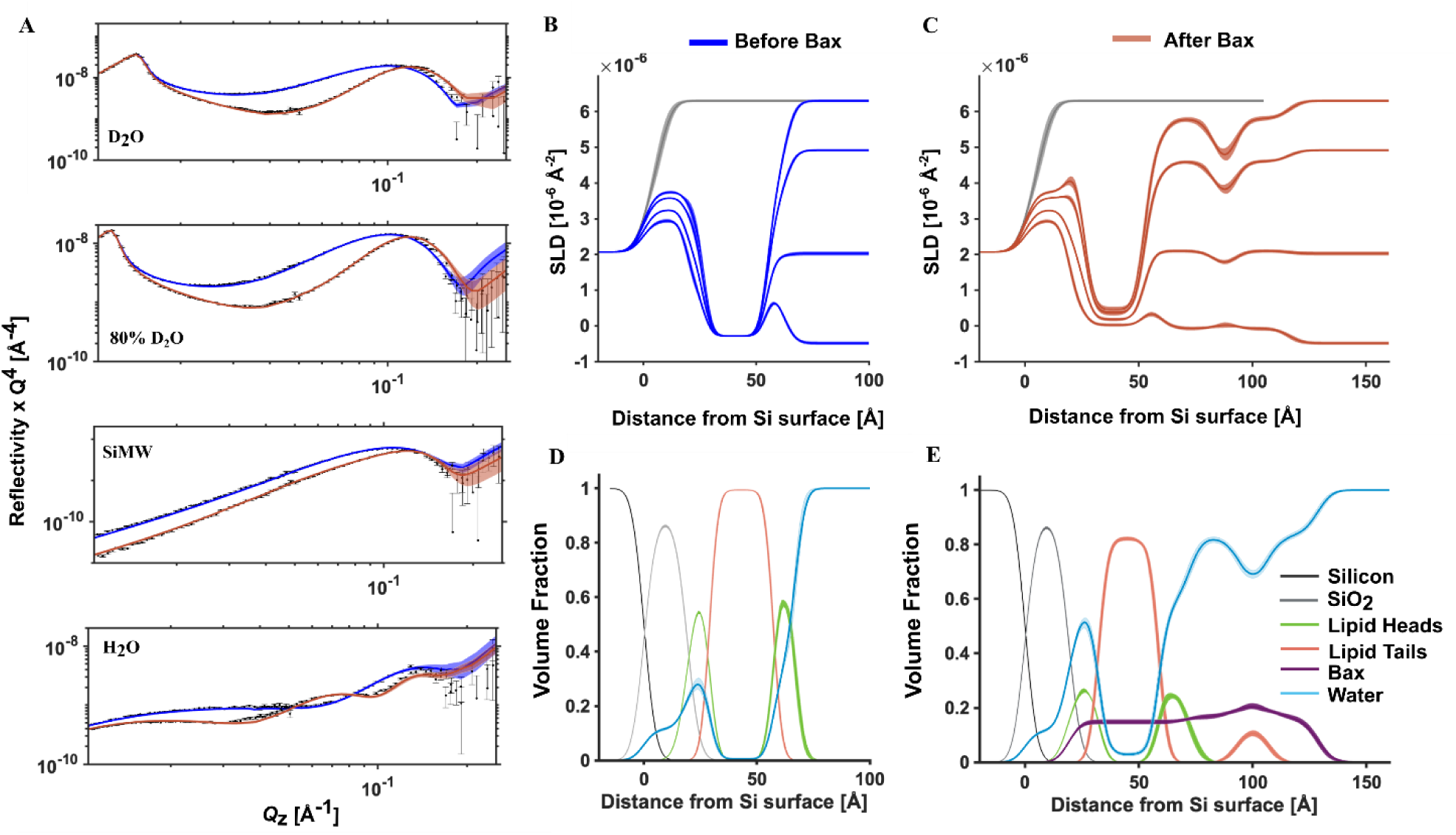
NR data showing Bax-induced pore formation in a POPC only bilayer. NR data (error bars) and model data fits (lines; see also supplement Table 1) from a natural-abundance hydrogen POPC (h-POPC) SLB before (blue) and after (red) the interaction of (h-)Bax are shown in four differing solution isotopic contrast conditions being D_2_O, 80% D_2_O, silicon-matched water (Si-MW) and H_2_O buffer solutions (A). The scattering length density (SLD) profiles are shown for the surface structure before (blue, B) and after (red, C) the h-Bax interaction, the bare substrate SLD profile is also shown (grey, B and C). The corresponding component volume fraction profiles are shown before (D) and after (E) the h-Bax interaction as determined from the NR fits. Individual components are color-coded as indicated, with the Bax protein distribution in purple. Note that after the interaction of the protein there is a lower lipid content in the SLB and a new distribution of lipid on the membrane surface. Line widths in the NR data fits represent the 65% confidence interval of the range of acceptable fits determined from Monte-Carlo-Markov Chain (MCMC) error analysis and the line widths in the SLD and volume fraction profiles represent the ambiguity in the resolved interfacial structure determined from these.

**Table 1:**
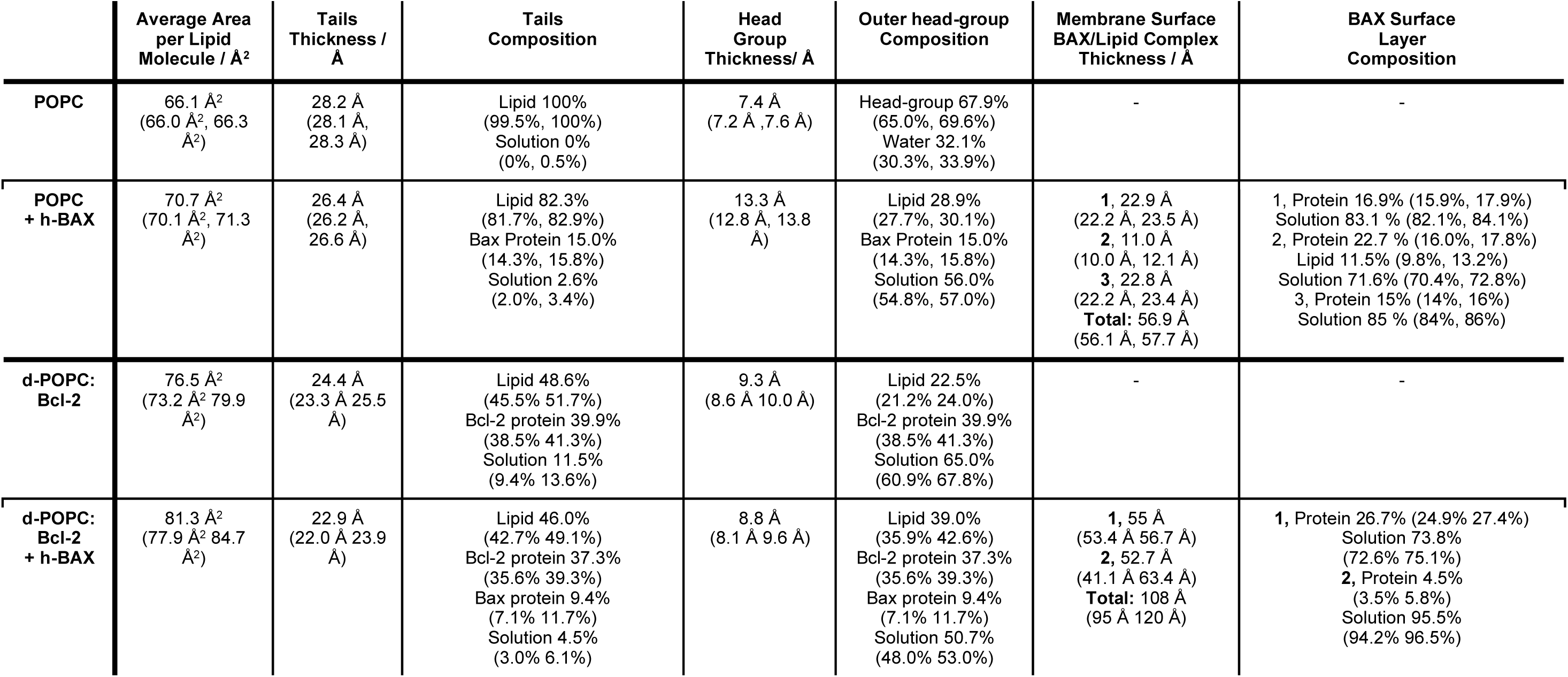
The resolved structural components before and after the interaction of Bax with SLB composed of POPC and POPC containing Bcl-2 protein. Bax Surface layer composition described as layers numbered outwards from the membrane, where 1 is membrane adjacent. *Values in parentheses represent the 65% confidence intervals determined from MCMC resampling of the experimental data fits.

As summarized in Table 1, the POPC only bilayers displayed a surface coverage of nearly 100 ±1 % lipid, a value reduced to 82 ± 1 % upon addition of h-Bax. The water content in the membrane hydrophobic core increased from 0 to 2.6 ± 0.7 volume %, reflecting the Bax induced membrane perforation. Even the bilayer thickness decreased by ca. 2 Å, a similar change as observed for CL containing bilayers upon Bax induced pore formation (*14*). The obtained volume fraction values for Bax protein and water in the POPC bilayers support significant membrane perforation by Bax similar as observed with CL containing bilayers which are closer to MOM-like systems (*14, 29, 30*). While in those membranes CL drives activation of Bax into pore forming oligomers in a faster CL dependent way, here in POPC this activation occurs on a slower hourly time scale, with a time constant of Bax-binding of 739 +/− 22 min, see ATR-FTIR analysis in Fig S6 and Fig S8, vs 175 +/− 15 min for 10% CL bilayers (*14*). However, the Bax mediated pore formation mechanism appears consistent This clearly shows that direct activation of Bax by lipids alone does not rely on the presence of anionic lipids (*27, 31, 32*) but can take place on neutral bilayers containing zwitterionic lipids, albeit on a slower time scale of several hours. This activation must include the conversion of inactive soluble Bax monomers to membrane-associated active Bax monomers (*2, 11, 31, 33*). The interaction with the lipids then drives the conversion of the Bax via its α2-α5 core domain into active dimers causing further assembly into 6-10 monomer large subunits and into pores (*30, 34, 35*), and finally releasing apoptotic factors, causing irreversible cell death. Assembly of additional Bax into multimeric states can further increase pore size by a kind of organized clustering near membrane-rupturing pores (*14, 15, 36*).

### Bax binds to Bcl-2 and oligomerizes on the membrane surface preventing pore formation

Bcl-2 containing POPC vesicles were deposited onto silicon supports as SLBs to form composite protein-lipid models of the MOM. As seen in the NR data and corresponding data analysis in Fig 2, Bcl-2 was found to be located within the lipid bilayer of the MOM model in agreement with our previous observations on similar systems (*22, 37*). Electron microscopy (EM) imaging of the protein-lipid vesicles used to fabricate the planar MOM models supported this analysis showing the presence of Bcl-2 within the lipid bilayer (See Fig 2 F).

**Figure 2.**
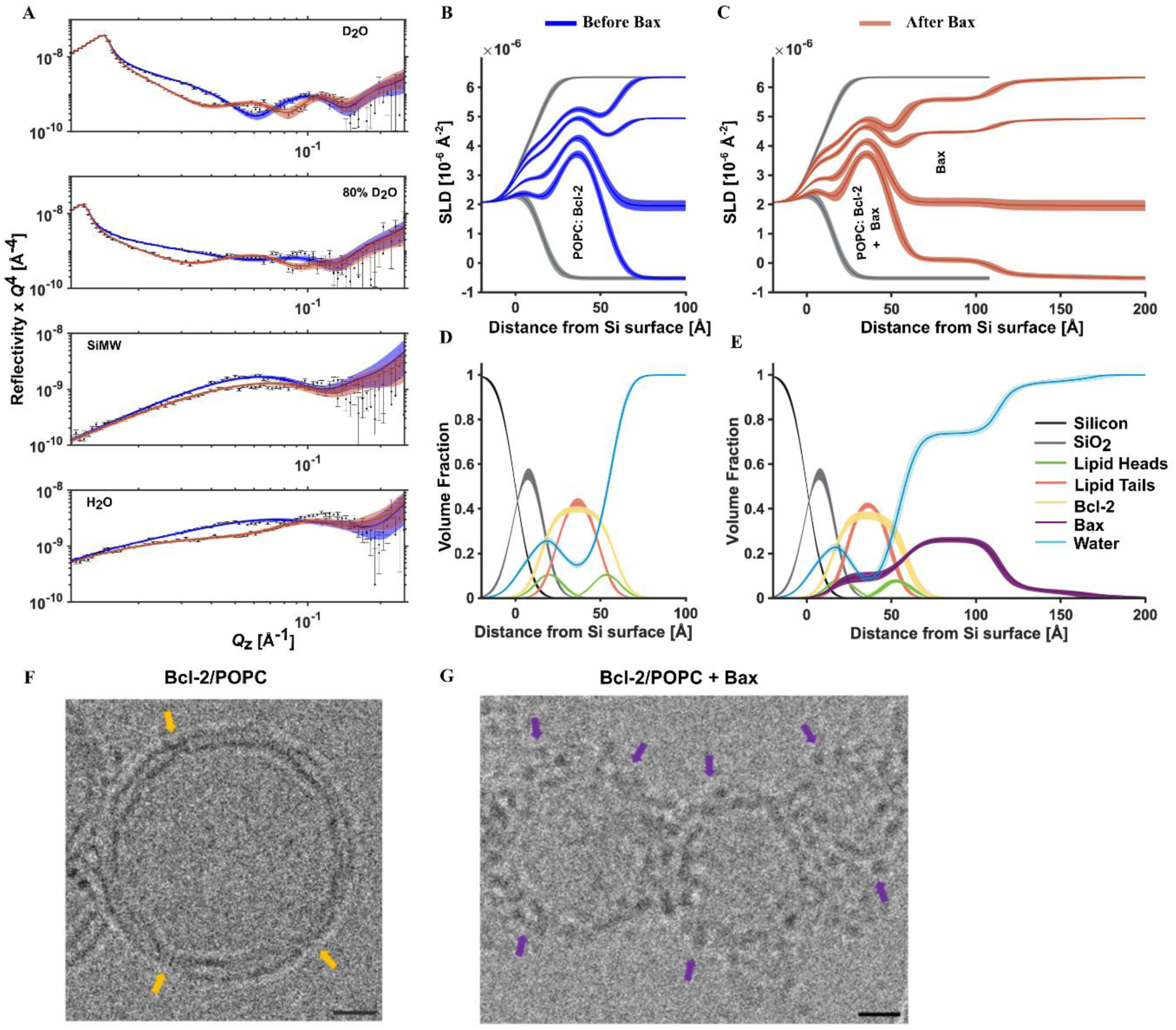
Bax binds to Bcl-2 containing membrane without lipid removal. NR data (error bars) and model data fits (lines; see also supplemental Table 1) from a d-POPC/Bcl-2 SLB before (blue) and after (red) the interaction of natural abundance hydrogen (h-)Bax are shown in four differing solution isotopic contrast conditions being D_2_O, 80% D_2_O, Si-MW and H_2_O (A) buffer solutions. The scattering length density (SLD) profiles are shown for the surface structure before (B) and after the h-Bax interaction (C). The corresponding component volume fraction profiles are shown before (D) and after (E) the h-Bax interaction as determined from the NR fits. Individual components are color-coded as indicated, with the Bcl-2 distribution in orange and the Bax protein distribution in purple. Line widths in the NR data fits represent the 65% confidence interval of the range of acceptable fits determined from Monte-Carlo-Markov Chain (MCMC) error analysis and the line widths in the SLD and volume fraction profiles represent the ambiguity in the resolved interfacial structure determined from these. Complementary Cryo-EM images of Bcl-2/POPC vesicles are shown indicating the presence of Bcl-2 (orange arrows) within the lipid bilayer (F) and the binding of discreet Bax distributions (purple arrows) onto the vesicular surface without disruption (G), consistent with the NR findings. The scale bar shown in 10 nm.

In the absence of Bcl-2, Bax interaction with POPC membranes led to membrane disruption by pore formation and the transfer of lipids into protein-lipid complexes on the bilayer surface (*38*). In comparison the Bax interaction with the POPC/Bcl-2 MOM models from repeated measurements (see Fig 2, 4 and SI Fig S1 and S2) only led to the presence of distribution of Bax across the bilayer depth and on its surface, and in most cases a minor bilayer thickening (∼2 Å, see Table S3), likely due to a change in tilt angle of the bilayer lipids needed to accommodate Bax penetration into the film. Together these results show unambiguously that the presence of the anti-apoptotic Bcl-2 changes the nature of the Bax interaction with the MOM surfaces compared to lipid only MOM models.

The amount of Bax on the membrane surface showed a direct correlation with the Bcl-2 content of the MOM models prepared (Fig 4 A-C and Table 2). This suggested a direct interaction between the two proteins and the likely formation of Bcl-2/Bax complexes at the membrane interface, consistent with previous observations (*37, 39*). The very clear change in the distribution of Bax away from the membrane interior to the surface, suggests that Bax is sequestered into these complexes by Bcl-2, thus preventing it from interacting with the lipids to form apoptotic pores.

**Table 2.** A comparison of bound Bax to Bcl-2 content in the MOM models studied by NR where no membrane disruption was observed. MOM bound Bax layers of similar coverage within the total membrane surface protein envelope were assigned as the distinct Bax distributions described here.

| Lipid composition | POPC |  |  | 9:1 (mol/mol) POPC:CL |  |
| --- | --- | --- | --- | --- | --- |
| Bcl-2 volume coverage in bilayer/% | 9.9<br>(9.5 10.1) | 14.2<br>(12.3 16.2) | 39.9<br>(38.5 41.3) | 24.5<br>(23.7 25.3) | 38.2<br>(37.4 39.0) |
| Bax surface bilayer proximal distribution thickness/Å | 88.4<br>(78.0 98.9) | 67.8<br>(62.7 73.8) | 55 Å<br>(53.4 56.7) | 52.6<br>(51.0 54.0) | 65.4<br>(64.0 66.7) |
| Bax surface bilayer proximal distribution coverage /% | 13.0<br>(11.9 14.0) | 8.0<br>(8.0 10.0) | 26.2<br>(24.9 27.4) | 21.2<br>(19.9 22.6) | 28.8<br>(28.2 30.1) |
| Bax surface bilayer distal distribution thickness/Å | 80.8<br>(72.6 87.6) | n/a | 52.7 Å<br>(41.1 63.4) | n/a | n/a |
| Bax surface bilayer distal distribution coverage /% | 5.8<br>(4.9 6.7) | n/a | 4.5<br>(3.5 5.8) | n/a | n/a |
| Total membrane bound Bax layer thickness/Å | 163<br>(154 174) | 67.8<br>(62.7 73.8) | 108<br>(95 120) | 52.6<br>(51.0 54.0) | 65.4<br>(64.0 66.7) |

Bax is a 21 kDa globular protein with a maximum diameter in its solution folded form of ∼40 Å (*12, 38*) (see SI Fig S12). The length scales of the Bax distributions in the presence of Bcl-2 were in MOM models was, in all cases, significantly larger than this value (see Fig 2, 4 and Table 2). Increasing the Bcl-2 content on the membrane produced shorter distributions with more Bax at the MOM surfaces. The length scales of the Bax distributions in the presence of Bcl-2 suggest that Bax is present as layers corresponding to ∼2-4 vertical protein units, suggesting oligomerization (see Fig 4).

These tapering distributions were most clearly seen for Bax bound to the ∼40% Bcl-2 containing MOM model (Fig 2). Here a 20% coverage of Bax was found in a ∼55 Å membrane proximal distribution which decreased sharply to 6% for an additional ∼53 Å Bax distribution bound to this. The ∼55 Å plus ∼53 Å tapering distribution is suggestive of a high-coverage layer of Bcl-2 bound Bax dimers at the membrane/water interface onto which additional Bax dimers bind. Indeed, alpha-fold modelling of Bax dimer structures (see SI section 3) suggested an average diameter of ∼49 Å with a maximum length of 65 Å for these. The ∼55 Å and ∼53 Å distributions found here are in good agreement with these length scales.

Cryo-EM imaging of Bax binding to Bcl-2 containing POPC vesicles supported the NR derived findings. Prior to Bax binding Bcl-2 was observed embedded in the lipid bilayer as distinct “notches” in-line with previous observations by us for the trans-membrane region of the barrel assembly machinery (*38*) (see Fig 2 F). Upon the binding of Bax to the vesicle surface no disruption of the vesicles (*39*) was observed and discreet distributions of the Bax proteins on the membrane surface were observed matching the Bcl-2 bound Bax layer revealed by NR (see Fig 2 G).

### Kinetics of Bax/Bcl-2 complex formation

A combination of time resolved (TR-)NR and attenuated total reflection (ATR-)FTIR was used to examine the time-dependence of the process by which Bax oligomerizes on the Bcl-2 containing MOM models.

Analysis of the change in the increase in the protein amide I band as Bax accumulated on the surface of a Bcl-2 containing deuterated (d-)POPC bilayer revealed a two-stage binding process with an initial fast component with a time constant of 9 ± 1 minutes and a slower secondary process with a time constant of 148 ± 11 minutes (Fig 3 C and D). TR-NR analysis had a lower time resolution (∼15 minutes for TR-NR vs. 80 seconds for ATR-FTIR), however, the analysis of the change in membrane surface protein content uncovered the slower secondary process also found in the ATR-FTIR measurements (time constant of 163 ± 11 minutes, Fig 3 D, inset).

**Figure 3.**
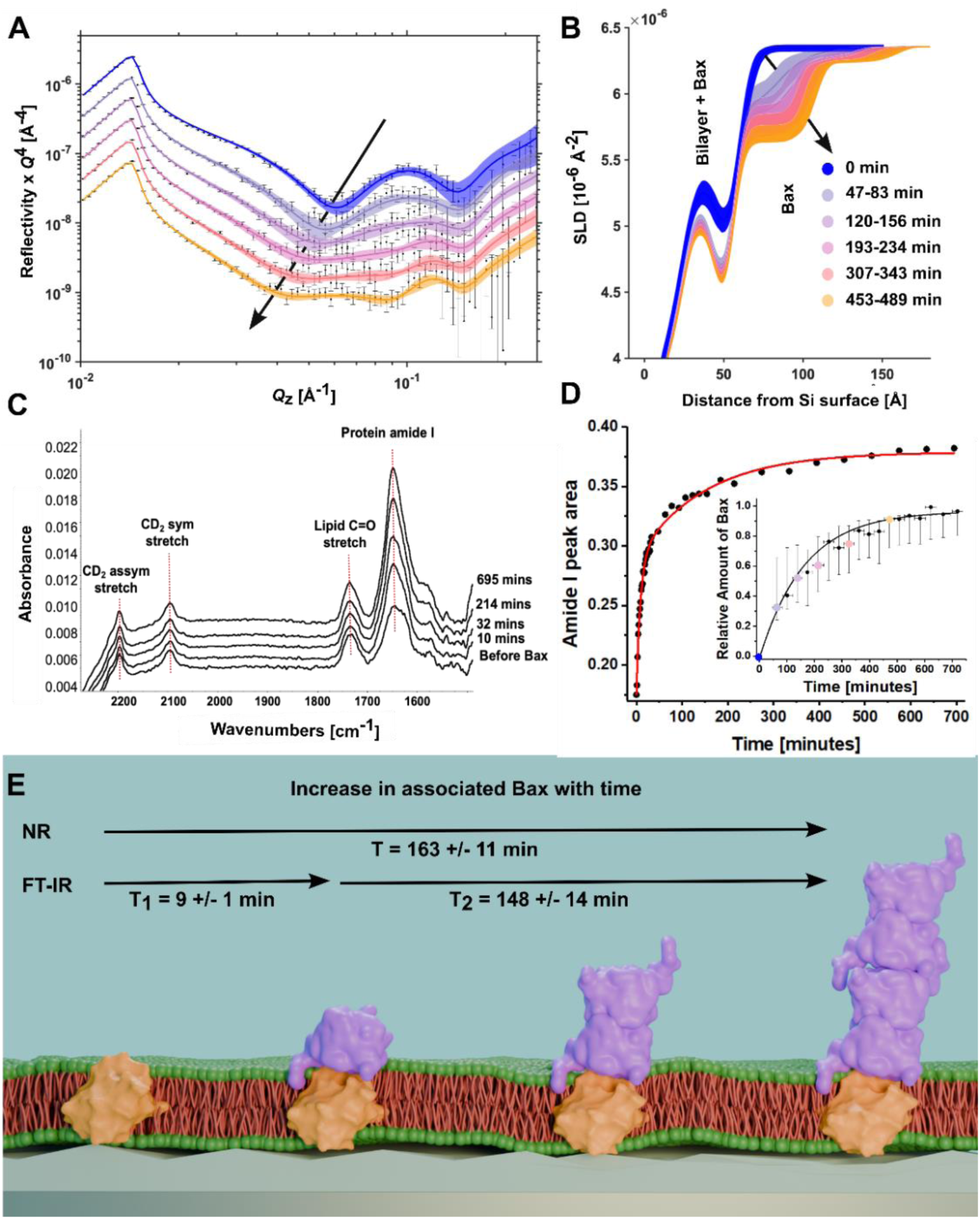
TR-NR and ATR-FTIR analysis of the kinetics of Bax binding a Bcl-2 containing d-POPC bilayer. TR-NR data (error bars) and model data fits (lines) in the D_2_O solution contrast obtained during Bax binding are shown (A) as are the SLD profiles from the MOM model surface/solution interface showing the increased accumulation of Bax on the SLB surface, black arrows indicate the change with time. (B) Complementary ATR-FTIR data is shown (C) depicting the increase in the Amide I during a Bax binding. Analysis of the increase in the Amide I peak vs. time (D) revealed a two-stage kinetic process with a fast (9 min) followed by a slower (148 min) process, see SI Figs S7 and S8. Inset kinetic analysis of NR data showing increase in Bax volume fraction with time, see S.I. Table S4 for detail. Each point represents a different NR dataset, coloured points match data sets shown in A and B. Analysis of the increase in the protein on the membrane surface from TR-NR analysis revealed initial binding of Bax monomers followed by dimerization then oligomerisation onto this, a schematic representation of this process is given (E), with protein depictions based on protein crystal structures 1F16 (Bax (*11*)) and IG5M (Bcl-2 (*28*)).

TR-NR analysis revealed the structural changes taking place on the MOM model surface during these processes. Initially a monomeric layer of Bax binds from solution onto the Bcl-2 containing MOM model. This binding was likely the short (T1) process revealed by ATR-FTIR. From this initial point the thickness of the Bax layer doubles to a ∼50 Å layer consistent with a Bax dimer which increases in membrane surface coverage and finally an additional Bax dimer-like layer appears bound to this which also increases in surface coverage (See Fig 3 B and a schematic interpretation of this in E). The oligomerization of Bax on the membrane surface following monomer binding to Bcl-2 is interpreted to be the slow secondary binding process revealed by both ATR-FTIR and TR-NR.

### Cardiolipin aids Bcl-2 in anchoring Bax to the membrane surface

The mitochondria-specific lipid CL plays a key role during apoptosis in facilitating recruitment of Bax to the MOM, the subsequent insertion of Bax into the membrane and its final perforation (*14, 29, 31, 40, 41*). The presence of CL accelerates and amplifies the Bax perforation significantly (*8*) compared to the behavior seen for the CL-free POPC systems used here (Fig 1 and 5). Nevertheless, the presence of Bcl-2 in CL containing membranes sequesters Bax in a very similar way as observed by us using POPC only bilayers, as the comparison of steady-state NR results for various Bcl-2 levels in POPC versus POPC/CL (9:1 molar ratio) membranes clearly show (Fig 4).

**Figure 4.**
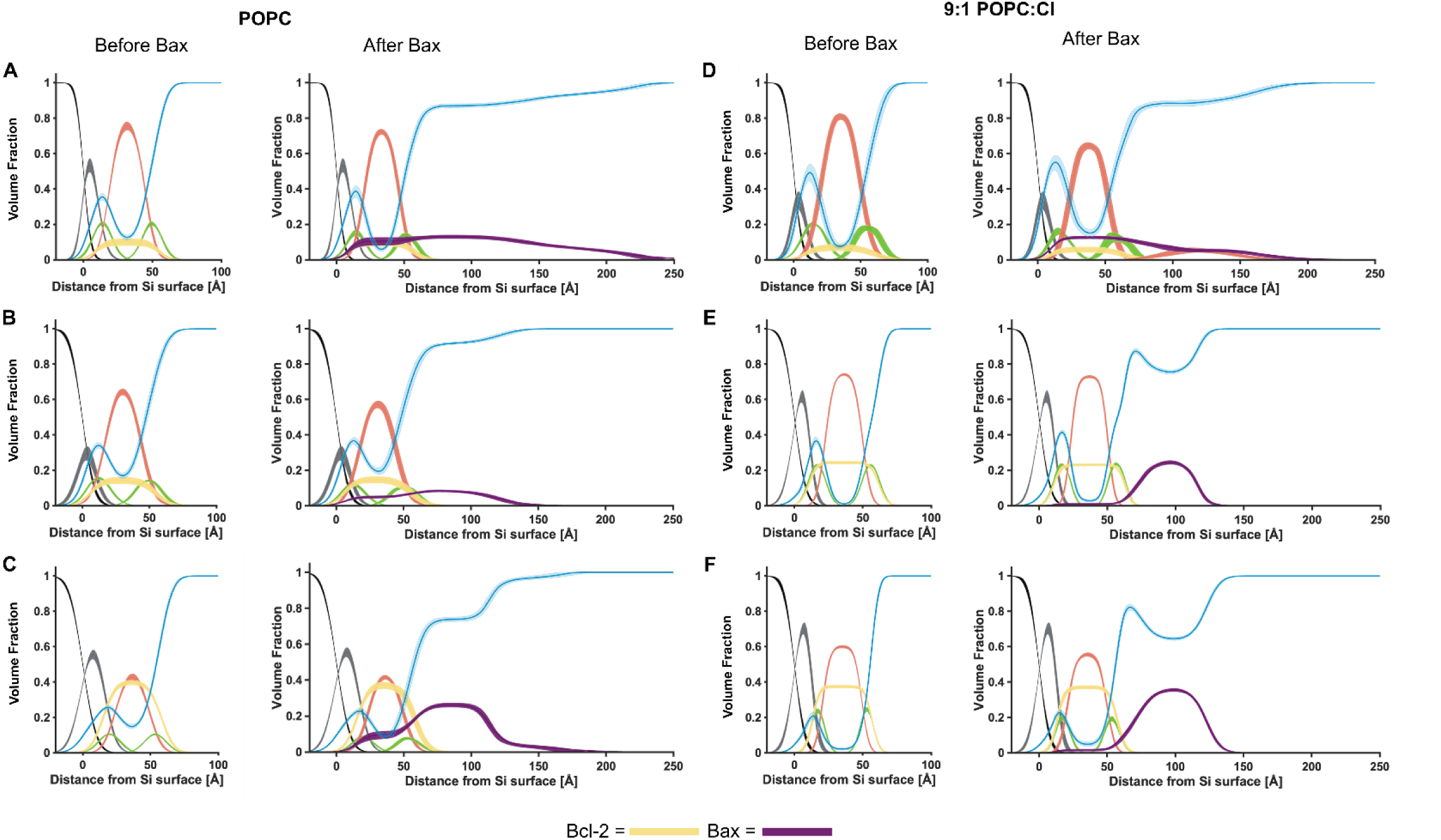
The positive correlation between bound Bax and SLB Bcl-2 content. Interaction of Bax with lipid bilayers containing an increasing volume fraction of Bcl-2, in POPC bilayers (A,B,C) and 9:1 POPC:CL bilayers (D, E, F) SLBs. Component volume fraction profiles determined from NR data analysis are shown before (left) and after (right) the interaction of Bax with the MOM models. Individual components are color-coded as indicated in Figures 1 and 2, with the Bcl-2 distribution in orange and the Bax protein distribution in purple. Line widths in the volume fraction profiles represent the ambiguity in the resolved interfacial structure determined from the 65% confidence interval of the range of acceptable fits from Monte-Carlo-Markov Chain (MCMC) error analysis of the NR data. NR data sets, model data fits and SLD profiles used to determine these volume fraction profiles are shown in Fig 2 and SI Figs S1-S5.

**Figure 5.**
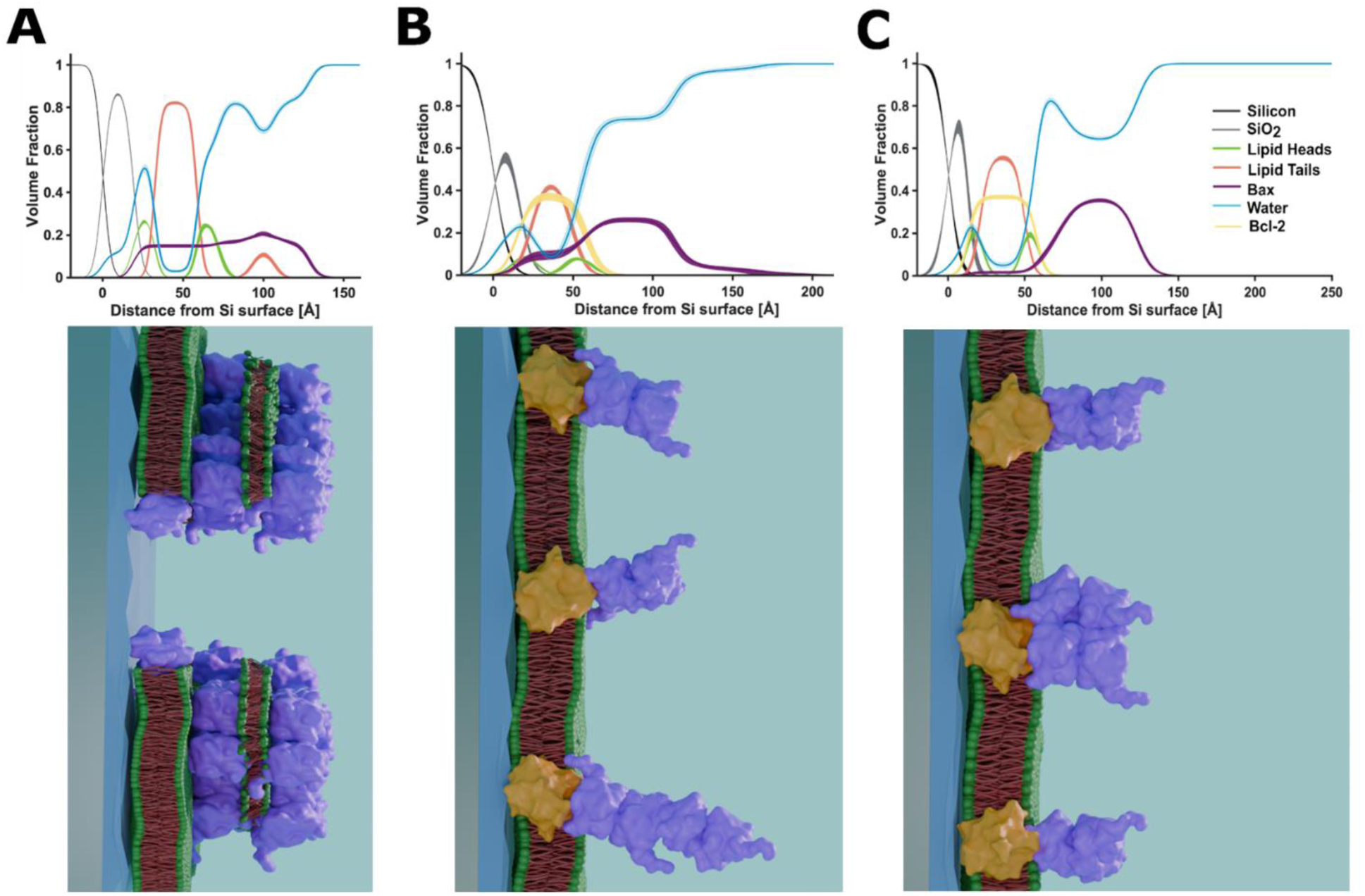
A comparison of NR derived component volume fraction distributions from Bax binding to MOM models with and without Bcl-2 and schematic representations of the derived distributions of components across the protein-lipid membranes. Bax binding to a POPC only bilayer (A) was associated with MOM lipid redistribution and increased water content indicative of pore formation whereas Bax binding to Bcl-2 containing MOM models with POPC (B) or POPC/CL (C) was associated with Bax membrane surface binding without membrane disruption.

In MOM models with both the pro-apoptotic CL (10% mol/mol lipid) and a relatively low amount of the anti-apoptotic Bcl-2 present (6% volume fraction) in the SLB there appeared to be competition between the Bcl-2 and CL for Bax leading to a mixture of Bax-Bcl-2 complexes and Bax lipid clusters (following poration) on the bilayer surface (Fig 4 D and Table 2). However, with a larger Bcl-2 volume fraction within the membrane Bax binding appeared to bind preferentially with Bcl-2 given that only Bax oligomers only found on the SLB surface (Fig 4, E and F). As with the Bcl-2/POPC SLBs there was a direct correlation between the Bcl-2 SLB content and the volume fraction of Bax oligomers on the surface (Fig 4, E and F and Table 2). Indeed a ∼29% Bcl-2 content within the SLB produced a ∼20% bound Bax coverage and a ∼40% Bcl-2 content produced a ∼30% Bax coverage on the membrane surface. These Bax layers were both thinner and had a higher coverage than the equivalent Bax oligomers found on the POPC/Bcl-2 SLB surfaces being ∼50-65 Å in total thickness (53 and 65 Å across two independent measurements, see Table 2) which is suggestive of a Bax dimer like structure (see SI section 3). This observation, combined with the lack of a membrane embedded Bax distribution is suggestive of CL playing a role in Bcl-2/Bax complex formation across the MOM which differs from that of POPC only, possibly forcing Bax oligomerization across the plane of the membrane rather than away from its surface as was observed for the Bcl-2/POPC models.

### Discussion

The observed sequestration of Bax by membrane-embedded Bcl-2 in the model membrane systems described here reflects the key features of the proposed Bcl-2 action *in vivo*, such as in Bcl-2 overexpressing cancer cells, namely the inhibition of Bax from initiating apoptotic cell-death (*2, 6, 23, 42*).

The hetero-dimerisation of Bcl-2 and Bax, has been long hypothesized as the key apoptotic blocking mechanism and has been largely investigated through co-precipitation, protein binding assays and cell-based studies (*42, 43*). It has been previously shown that the Bcl-2 protein contains a hydrophobic protein core with its BH3-BH1-BH2 region forming an extended groove interface, which is thought to recognize apoptotic proteins like Bax via their specific Bcl-2 Homology 3 (BH3) death motifs (*28, 42, 44*). These Bax motifs are known to contain a series of hydrophobic residues and engage with the Bcl-2 with nM affinity (*44*), and enable Bax to sequester into a Bcl-2/Bax complex with similar affinity *in vitro* (*37, 45*) as *in vivo* (*46*).

In the TR-NR and FTIR experiments described here (Fig 3), an initial formation of a Bcl-2/Bax complex across the membrane was observed, consistent with a Bcl-2/Bax hetero-dimer in its distribution on the membrane. This happens on a timescale of 9 min upon a fast initial and independent process of Bax association to the membrane. The complex is consistent with a 1:1 complex, also observed in intact mitochondria between Bax and Bcl-x_L_ (*47*) a close relative to Bcl-2, and we expected the same for our system with full-length human, membrane embedded Bcl-2. The 1:1 complex is also consistent with our previous basic affinity studies (*23, 37*). Our results suggest that initially Bcl-2 proteins sequester Bax (Table S2 and Fig 3) likely through membrane-mediated conversion of inactive Bax monomers into their active monomeric and dimeric forms (*2, 11, 31, 33*), preventing the onset of pore creation. Bcl-2 is thought to recognize Bax via its exposed BH3 motif as shown previously (*23, 42*); and the interaction occurs at membrane level near the lipid interface region as depicted in Fig 3E.

However, we also observe a slower second step in the kinetics of Bax binding to the Bcl-2 containing MOM models. NR characterization of the final steady state structures showed a Bax distribution consistent with Bax layers corresponding to 2-4 monomeric units in length from the membrane (and Bcl-2) surface, as illustrated in Figure 5. Bax-Bax oligomerization has been characterized previously as active and in-active cytosolic Bax dimers (*11, 13, 33, 48*) and is further observed in its pore-formation in the absence of Bcl-2 (*12, 30, 34, 35*). To reveal Bax’s BH3 hydrophobic domain, required either for membrane perforation, self-dimerization or likely Bcl-2/Bax hetero-dimerization, it must undergo a conformational change which as well as freeing the BH3 domain (*2*), exposes the Bax canonical groove within its core domain (α1-α5 helices). This can act as an interaction site for further Bax oligomerization (*49*). The Bax distribution across our repeat measurements strongly suggests oligomeric Bax binding indicative of a mechanism by which an initially Bcl-2-bound Bax layer binds additional Bax, inhibiting any free Bax from membrane perforation.

However, we cannot completely differentiate if Bcl-2 initially interacts only with Bax monomers, and additional Bax is subsequently assembled with the Bcl-2 bound Bax into a dimer, or if Bcl-2 directly can also recognize active, partially penetrated Bax dimers at the membrane level. Nevertheless, as deduced from our kinetic structural data, the temporal development of localized Bax clusters near Bcl-2 (as shown in Fig 3). Most importantly, despite the vast excess of Bax proteins at the surface there is no sign of any membrane damage demonstrated by both NR analysis and EM imaging (Fig 3 E).

The anionic CL is an important driving force in Bax-induced membrane pore formation, significantly increasing perforation activity and kinetics compared to CL-free neutral lipid bilayers (*14, 27, 32, 41*). In the presence of Bcl-2 the influence of CL on the Bax interplay with Bcl-2 is much less visible but still leads to a difference in observed clustering of Bax at the membrane near the Bcl-2 (Fig 5). Clearly, pore formation is still inhibited unless if the Bcl-2 abundance is very low, as seen in Fig 4D. Since CL has anionic headgroups its affinity for Bax is higher than that of POPC. Therefore, upon Bcl-2 binding the Bax redistribution is close to this highly charged membrane region with nearly no Bax deeply inserted into the membrane hydrophobic core and less Bax clustering further away from the membrane surface as seen for CL-free membranes (Fig S4). Since the main features for sequestering Bax by Bcl-2 are similar for both membrane systems, we suggest the main binding interface between Bcl-2 and Bax must be located near the bilayer interface.

### Conclusions

The presence of the anti-apoptotic Bcl-2 prevents Bax induced pore formation by sequestering Bax into Bax/Bcl-2 complexes. Onto these complexes Bax oligomerises and these events inhibit the initial stages of intrinsic apoptosis even in the presence of pro-apoptotic CL. Only at very low Bcl-2 volume fractions and in the presence of CL was Bax able to overcome Bcl-2 to form pores in the model systems studied here. Overexpression of Bcl-2 has been found in a number of cancers where it contributes to disease proliferation by reducing apoptosis. To our knowledge this study represents the first direct molecular level structural evidence obtained in a membrane of the role of the Bcl-2/Bax complexation in preventing apoptosis uniquely demonstrating that Bax sequestration is not only due to its binding to Bcl-2 but also subsequent binding to itself.

Our findings provide a structural and molecular basis for understanding the cell-protecting mechanism of anti-apoptotic members of the Bcl-2 family. A mechanism that allows cancer cells to escape the apoptotic drive from oncogenic stress and cytotoxic therapies, and opens up new avenues for innovative cancer drugs targeting Bcl-2’s overprotective functioning.

## Materials and Methods

### Materials

#### Lipids

Tail-deuterated 1-palmitoyl-2-oleoyl-d63-glycero-3-phosphocholine (d-POPC) was synthesized by the Deuteration and Macromolecular Crystallography (DEMAX) platform at the European Spallation Source (ESS), Lund, Sweden using the published method (*50*). Natural abundance hydrogen 1-palmitoyl-2-oleoyl-glycero-3-phosphocholine (h-POPC) and Cardiolipin from bovine heart (CL) were purchased from Sigma Aldrich as solid powders.

##### Expression and purification of protonated (h-Bax) and deuterated (d-Bax) proteins

Production of Bax protein for NR, EM and ATR-FTIR studies was accomplished by following the previously published method (*51*). Deuteration (> 90%) of Bax was carried out in a similar fashion, but using a M9 minimal media recipe as: 1 liter media was prepared by mixing 13 g KH_2_PO_4_, 10 g K_2_HPO_4_, 9 g Na_2_HPO_4_, 2.4 g K_2_SO_4_, 2 g NH_4_Cl, 2.5 ml of MgCl_2_ (2.5 M stock), 1 ml thiamine (30 mg/ml stock), 2 g glucose (non-deuterated), and 2 g NH_4_Cl (non-deuterated), followed by addition of trace elements (1 ml of 50 mM FeCl_3_, 20 mM CaCl_2_, 10 mM MnCl_2_, 10 mM ZnSO_4_, 2 mM CoCl_2_, 2 mM CuCl_2_, 2 mM NiCl_2_, 2 mM Na_2_MoO_4_ and 2 mM H_3_BO_3_), 100 μg/ml carbencillin and 34 μg/ml chloramphenicol. The pH of the media was adjusted to 6.9 and sterile-filtered before use.

##### Expression and purification of protonated Bcl-2

was carried out as previously reported (*52*). Reconstitution of Bcl-2 into proteoliposomes in POPC (1-palmitoyl-2-oleoyl-*sn*-glycero-3-phosphocholine) and cardiolipin (1,3-bis(sn-3’-phosphatidyl)-*sn*-glycerol) was accomplished by following the method as described (*51*). Successful incorporation of Bcl-2 protein into the various bilayer systems was verified by SDS-PAGE prior to the neutron reflectometry experiments.

### Methods

#### Neutron Reflectometry Measurements

NR measurements were performed on the white beam SURF and OFFSPEC (*53*) reflectometers at the ISIS Neutron and Muon Source (Rutherford Appleton Laboratory, Oxfordshire, UK) and on the Figaro reflectometer at the Institut Laue Langevin (ILL, Grenoble France), which use neutron wavelengths from 0.5 to 7 Å, 1 to 14 Å and 2 to 20 Å respectively. The reflected intensity was measured at glancing angles of 0.35°, 0.65°, and 1.5° for SURF, 0.7° and 2.0° for OFFSPEC and 0.7° and 2.3° for Figaro. Reflectivity was measured as a function of the wave vector transfer, Q_z_ (Q_z_ = (4π sin θ)/λ where λ is wavelength and θ is the incident angle). Data was obtained at a nominal resolution (dQ/Q) of 3.5% at ISIS, and 7.0% at ILL. The total illuminated sample length was ∼60 mm on all instruments. Measurement times for a single reflectometry data set (∼0.01 to 0.3 Å^−1^) was 40 to 180 minutes at ISIS, 20 to 60 minutes at ILL. Data collection times for kinetic data sets varied and can be seen as the x-error bar on Fig 3D (inset) and Fig 4D.

Details of the solid-liquid flow cell and liquid exchange setup used in the experiments described here are reported by us previously (*54*). Briefly, solid liquid flow cells containing piranha acid (sulphuric acid, hydrogen peroxide and water mixture) cleaned 111 silicon substrates (15 mm × 50 mm × 80 mm with one 50 mm × 80 mm surface polished to 3 Å root mean squared roughness) were placed onto the instrument sample position and connected to instrument controlled HPLC pumps (Knauer Smartline 1000) which enabled programmable control of the change of solution isotopic contrast in the flow cell.

##### Vesicle preparation for NR

POPC vesicles were prepared for deposition by hydrating the lipid in D_2_O to a concentration of 0.2 mg ml^−1^, bath sonicating for 30 minutes and tip sonicating on ice for 10 minutes (1s on 2s off) to produce vesicles of roughly 100 nm in diameter. Bcl-2 containing vesicles were prepared by hydrating the pellet in deposition buffer (10 mM Sodium Citrate pD/H 3.8 25 mM NaCl 1.25 mM CaCl_2_), centrifuging back to a pellet whilst discarding the supernatant and resuspending the pellet into deposition buffer to a lipid concentration of 0.2 mg ml^−1^. The Bcl-2 containing vesicles were then tip-sonicated on ice for 10 minutes (1s on 2s off), to an average diameter of roughly 150-200 nm, ensuring minimal time between sonication and injection into the ATR-FTIR or NR flow cell.

###### Lipid Membrane Deposition

Initially, the clean silicon substrates were characterized by NR in D_2_O and H_2_O buffer solutions. Then freshly sonicated lipid vesicle solutions (0.2 mg/ml) were introduced into the cells in the experiment deposition buffer solution of 10 mM Sodium Citrate pD/H 3.8 25 mM NaCl 1.25 mM CaCl_2_ and the samples incubated at 30±1°C for ∼1 hour before the non-surface bound vesicles were removed by flushing the cells with 15 ml (∼5 cell volumes) of the same buffer solution before a solution of pure D_2_O was flushed into the cell. This formed high-quality (i.e. high-coverage) supported lipid bilayers (SLBs) at the solid/liquid interface. The resulting bilayers were characterized by NR in the experiment buffer (20 mM Sodium Phosphate, pH 7.4, 50 mM NaCl, 1mM EDTA) under four solution isotopic contrast conditions being D_2_O, 80% D_2_O: 20% H_2_O, Silicon Matched Water (Si-MW, 38% D_2_O) and H_2_O buffer solutions. The remainder of the beamtime was conducted in this buffer.

###### Bax Interaction

Once characterisation of the SLB was complete the samples surface was placed in the correct experiment buffer isotopic contrast (D_2_O for h-proteins and H_2_O for d-proteins) and ∼6 ml of a 0.1 mg/ml Bax solution was injected into the flow cell (the cell volume is 3 ml) either by hand (SURF) or using a syringe pump (OFFSPEC and INTER, AL1000-220, World Precision Instruments; Figaro, The Harvard Apparatus Pump 33 DDS). In most cases the interaction of the protein with the SLB was monitored by NR with data sets collected continuously until an equilibrium interaction between the protein and the SLB was verified by no further changes in the data being observed against time. At this point a final equilibrium data set was collected and then the excess protein was flushed from the cell and the structure of the surface protein-lipid complex was examined by NR under three solution isotopic contrast conditions (D_2_O, Si-MW and H_2_O). It should be noted no difference was found in any sample between the equilibrium Bax bound data before and after flushing of the excess protein suggesting the protein-lipid complexes formed at the sample surface were stable.

###### NR Data Analysis

NR data was analysed using the RasCal software (A. Hughes, ISIS Spallation Neutron Source, Rutherford Appleton Laboratory) which employs optical matrix formalism (*55*) to fit layered models of the structure across bulk interfaces and allows for the simultaneous analysis of multiple NR data sets collected under different sample and isotopic contrast conditions and permits them to be fully or partially constrained to the same surface structure in terms of thickness profile but vary in terms of neutron scattering length density. For additional details see SI section 1.

## Acknowledgments

GG acknowledges support from the national infrastructures SciLifeLab, SwedNMR and Knut and Alice Wallenberg foundation programme “NMRforLife”. The synthesis of d-POPC was carried out at the DEMAX Platform resulting from proposals CTU4H35A and 128669, at the European Spallation Source ERIC. The persistent identifiers for the samples are doi:10.5281/zenodo.14002732 and doi:10.5281/zenodo.4160419. Schematics were generated in Blender.

## Funding

GG, LAC and HPWK acknowledges financial support from:

Swedish Research Council 2021-00167, 2016-06963

Kempe Foundation JCK-1321

Umeå Insamlingsstiftelsen FS 2.1.6-2396-18

This work was supported by ISIS Neutron and Muon source beam time awards 2210172, 2010295 and 1919323 and Institut Laue-Langevin beamtime award 10.5291/ILL-DATA.8-02-999.

## Author contributions

S.A., G.G., L.A.C. and H.P.W.-K designed research; S.A., L.A.C., J.Å., S.K., N. P., H.P.W.-K., E.C.B., T.M.N. and G.G. performed research; A.E.L., O.B., J.-F.P. synthesized the deuterated lipid samples; S.A., L.A.C., H.P.W.-K., E. V. B., J. D. and G.G. analyzed data; S.A., L.A.C., G.G., J.Å., and H.P.W.-K wrote Manuscript.

## Data and materials availability

NR data and custom model scripts used in NR data fitting as well as ATR-FTIR data are available via https://doi.org/10.5281/zenodo.14698607. Information on the purification of tail deuterated POPC can be found at https://doi.org/10.5281/zenodo.4160419 and https://doi.org/10.5281/zenodo.14002732.

