## Supplementary Information for "Bcl-2 Oligomerizes Bax on the Mitochondrial Membrane Surface Preventing the Initial Stages of Apoptosis"

**Short title**: Formation of Bcl-2/Bax complex at membrane level prevents membrane poration.

Sophie E. Ayscough[1,2,3] Luke A. Clifton[1, *], Jörgen Ådén[4], Sebastian Köhler[5], Nicolò Paracini[2], James Doutch[1], Éilís C Bragginton[6], Anna. E. Leung[2], Oliver Bogojevic[2], Jia-Fei Poon[2], Tamás Milán Nagy[4], Hanna P. Wacklin-Knecht[2,3], Gerhard Gröbner[4, *]

[1] ISIS Pulsed Neutron and Muon Source, Science and Technology Facilities Council, Rutherford Appleton Laboratory, Harwell Science & Innovation Campus, Didcot, Oxfordshire, OX11 OQX, UK.

[2] European Spallation Source ERIC, ESS, P.O. Box 176, SE-22100 Lund, Sweden.

[3] Department of Chemistry, Division of Physical Chemistry, Lund University, P.O. Box 124, SE-22100 Lund, Sweden

[4] Department of Chemistry, University of Umeå, SE -901 87, Umeå, Sweden

[5] Lund Institute for Neutron and X-ray Scattering, Department of Chemistry, Lund University, P.O. Box 124, SE-22100 Lund, Sweden

[6] Electron Bio-Imaging Centre (eBIC), Diamond Light Source Ltd, Diamond House, Harwell Science and Innovation Campus, OX11 0DE, UK.

* Luke A. Clifton: *Gerhard Gröbner**:**

### Section 1: Additional Methods

#### Neutron Reflectometry Data Analysis:

Neutron Reflectivity data was fitted with the Rascal 2.0 package (*1*), using the custom model option whereby the fitted parameters defining the interfacial layers are given definitions and constraints. The structure of the substrates, the bilayers and the bilayer-protein interactions were determined by finding the parameters that minimized the chi2 value between the reflectivity data and the fitting model. The bilayers were fitted as 5-layer model, from the substrate to the subphase; a thin $SiO_2$ layer, an inner lipid head group layer, an inner tail group layer, an outer tail layer and an outer head group layer. The molar ratio of head to tail groups of the lipids were maintained by fitting a lipid area per molecule parameter, thickness of the bilayer head $t_{hg}$ and tail regions $t_t$ are given by:

$$t_{hg} = \frac{V_{hg} + n_w V_w}{APM}$$

and

$$t_t = \frac{V_t}{APM},$$

Where $V_{hg}$ , $V_t$ and $V_w$ are the average volumes of the head and tails groups and that of a water molecule respectively, $n_w$ is the number of water molecules per lipid head and $APM$ is the lipid area per molecule. In all the fits of this publication the APM and $n_w$ are constrained to be the same for both bilayer leaflets. The scattering length density of a component is defined by:

$$SLD = \frac{\sum b}{V},$$

Where $\sum b$ is the sum of the component scattering lengths and $V$ is the molecular volume of the component. The molecular volumes and scattering length densities of the lipid heads and tails were calculated from literature values (*2*), addition of their sub-component volumes and addition of their -component scattering lengths, full detail in the NR scripts included in our electronic supplementary information. The SLD of each layer is calculated by fitting the volume fraction of each component in that layer. In addition to the water per lipid heads parameter allowing for a volume fraction of water in the more hydrophilic head layer, we have a hydration parameter across the entire bilayer, both head and tail layers, to allow for any patchiness of the bilayers. For the bilayers containing the integral membrane protein Bcl-2, a single volume fraction (VF) of Bcl-2 is fitted across the bilayer layers. Each of the interfaces between layers in the model have an associated roughness, we allow a separate roughness to be fitted for the Si-$SiO_2$ interface $SiO_2$-bilayer interface, otherwise the bilayer interface roughness's are constrained to be the same fitted value.-bilayer interface, otherwise the bilayer interface roughness's are constrained to be the same fitted value.

For Bcl-2 containing bilayers on equilibrium binding of Bax, with exception of the POPC:CL bilayer with low Bcl-2 VF of about 5%, a seven-layer model was found to be the most reasonable model that satisfactorily resolved the features in the data. The first 5 layers were the thin $SiO_2$ layer and bilayer, the APM, for which the hydration and water per lipid head parameters were re-fitted after the Bax interaction but the Bcl-2 volume fraction was assumed to not change and was constrained to be the same as before Bax interaction using:

$$VF_{Bcl-2_{ab}} = VF_{Bcl-2_{bb}} \times \left(\frac{t_{b_{bb}}}{t_{b_{ab}}}\right),$$

Where $VF_{Bcl-2_{bb}}$ is the volume fraction of Bcl-2 in the bilayer prior to Bax interaction, $t_b$i. s the bilayer thickness before ($bb$ ) and after ($ab$ ) Bax interaction. This is done as we assume the amount of Bcl-2 at the interface will not change. We also allow for a volume fraction of Bax across the bilayer, although there was significantly less Bax insertion than in the lipid only data sets. The two added layers are protein layers, where the volume fraction of Bax and the thickness of the layers are fitted.

For the equilibrium binding of Bax to the POPC bilayers without Bcl-2, a similar model was used as previously for POPC:CL bilayers (*2*). This model is an eight-layer model of a disrupted bilayer and a lipid protein complex. The first five layers ($SiO_2$ + lipid bilayer) of the structure were the same as prior to the Bax interaction, with an increase in protein and water content and a decrease in the lipid content. The three added layers are a Bax-protein only layer next to the bilayer, a mixed Bax-protein/lipid layer and an additional Bax-protein layer adjacent to the bulk solution. A similar model was used to fit the POPC:CL bilayer with a low volume fraction of Bcl-2 (5%), in which lipid removal was also observed. The Bcl-2 was constrained, but the bilayer disruption was modeled as a decrease in lipid and increase in water content and 3 additional layers on top of the bilayer consisting of a Bax-protein only layer next to the bilayer, a mixed Bax-protein/lipid layer and an additional Bax-protein layer adjacent to the bulk solution.

**Analysis of time-resolved (TR)-NR data**

The TR-NR data describing the Bax interaction with a d-POPC: Bcl-2 bilayer, was measured in a single subphase contrast ($D_2O$) after the injection of Bax into the NR flow cell. This data was batch-fitted. The parameters of the final equilibrium Bax-bilayer fit were used as a fixed constraint, allowing for only two parameters of the Bax membrane surface component to vary. The best-fit of the TR-NR datasets was found allowing for the value of the volume fraction of Bax and surface layer thickness to vary, the maximum volume fraction of Bax being that found in the equilibrium fit. The calculated volume fraction value was applied at the same value to the volume fraction of Bax inside the bilayer, in the first membrane associated Bax layer and then the second Bax layer. Similarly, the thickness value was applied to the first and second Bax layers. Table S.4 shows the fit values and how this relates to the thickness and volume fraction parameters in the model.

**NR Error Estimation and plotting**

Bayesian analysis was used to calculate the confidence intervals of the neutron reflectivity model to data fitting parameters and therefore provide the error estimation for our calculated structures. Bayesian analysis was done in Rascal using Monte-Carlo-Markov Chain (MCMC) and Delayed-Rejection Adaptive Metropolis (DRAM) algorithms routines to refit the data from the already chi2 minimized fit *(*3). The parameter uncertainties are determined from the posterior distributions as the 65% confidence interval and the uncertainties on the scattering length density and reflectivity plots were generated from 1000 random samples from the Markov chains. These chain samples are used to generate the line shading in the reflectivity, SLD and volume fraction plots whilst the darker lines represent the best fit lines.

**Component Volume Fraction Profiles**

Volume fraction profiles detailing the distribution of components across the solid/liquid interface before and after the equilibrium interaction of Bax were produced using a bespoke script. MCMC Bayesian error estimation results and the relationship between the fitting parameters and interfacial structure in the RasCal custom model were used to determine the distribution of each structural component in the volume fraction vs. distance profile. The volume fraction of an individual component was calculated in 1 Å increments across the solid/liquid interface (the silicon/silicon dioxide interface set as zero). The mean, lower, and upper 65% confidence interval bounds of each component distribution were determined for every 1 Å segment using the MCMC error estimation results or derived parameters; these confidence intervals were then used to produce a line width error region above and below the mean values. The water distribution was calculated as the remaining unoccupied volume for each 1 Å slice and summed across the interface with the appropriate error propagation.

#### CryoEM sample preparation and data collection

Two 100 µL aliquots of Bcl-2 containing POPC vesicles were prepared at a protein concentration of 0.5 mg $ml^{-1}$ by resuspending pellets in buffer (20 mM Sodium Phosphate, pH 7.4, 50 mM NaCl, 1mM EDTA) and tip sonicating to an average diameter of about 200 nm. To one aliquot, 100 µl of 0.1 mg $ml^{-1}$ Bax was added and the samples incubated at 37° C for 1 hour. They were then frozen within 30 minutes.

Four microliters of freshly prepared proteoliposomes at a Bcl-2 protein concentration of 0.5 mg/ml were deposited onto glow discharged R1.2/1.3 Cu 300 mesh holy carbon grids (Quantifoil) prior to vitrification. Grids were plunge frozen using a Virtobot Mark IV (Thermo Fisher Scientific) (Blot force 2, Blot time 3 sec, temperature 22 degrees) and stored in liquid nitrogen prior to imaging. Data was collected at the electron Bio-Imaging Centre (eBIC) using a Titan Krios microscope (Thermo Fisher Scientific) equipped with a field emission gun operating at 300 keV, a Falcon 4i direct electron detector with a Selectris X imaging filter (Thermo Fisher Scientific). Data was collected with EPU software (Thermo Fisher Scientific) at a magnification of x130 000 with a corresponding pixel size of 0.921 Å/pixel. The total dose applied to the sample was 40 e-/$Å^2$.

#### Attenuated Total Reflection Fourier Transform Infra-Red Spectroscopy (ATR-FTIR)

Trapezoidal silicon substrates for attenuated total reflection infrared spectroscopy were obtained from Crystran (Poole, UK). These substrates were made to fit into a Specac (Orpington, UK) liquids ATR accessory which was, itself, fitted into the sample cavity Thermo-Fisher iS50 Infra-red spectrometer (Waltham, MA, USA). The substrates have four polished faces (to ~6 Å root mean squared roughness) the largest being a 72 mm × 10 mm face which was used as the sample surface. The IR beam enters the substrate through a polished face at 45° relative to the sample surface, total internal reflectance of the IR beam inside the substrate gives rise to six evanescent waves on the sample surface. IR spectra were collected at a resolution of 4 $cm^{-1}$.

To reduce the influence of water vapour on the IR spectra the iS50 instrument and ATR mirror assembly accessory was continuously purged with dry air from a Peak scientific (Glasgow, UK) $CO_2$ and water removing air purge. The Specac liquid ATR cell was modified to fit Omni-fit tubing which was connected to a syringe pump (AL1000-220, World Precision Instruments, Hitchin, UK). The experimental $D_2O$ buffer solution (20 mM Sodium Phosphate, pD 7.4, 50 mM NaCl, 1mM EDTA) was injected into the ATR flow cell which was then heated to 30±1°C using a water bath (Julabo, Seelbach Germany). $D_2O$ solutions were used for all ATR-FTIR measurements as the $D_2O$ bending mode is lower (1215 $cm^{-1}$) compared to $H_2O$ (1645 $cm^{-1}$) meaning limited contamination of protein amide I region by spectral bands from water *(*4). After buffer flushing a background spectrum was collected. Spectra were then collected monitoring the removal of water vapor from the spectrometer and mirror assembly until a steady state was reached. A 2nd background measurement was then collected before deposition of the SLB.

SLB deposition was conducted using the same methodology as described for NR measurements and monitored through the appearance of $CH_3$ / $CD_3$ asymmetric, $CH_2$ / $CD_2$ asymmetric and $CH_2$ / $CD_2$ symmetric stretches from the lipid tails at ~2950 $cm^{-1}$, 2920 $cm^{-1}$ and 2850 $cm^{-1}$ respectively and the appearance of a lipid carbonyl stretch at 1730 $cm^{-1}$. In the case of the d-POPC: h-Bcl2 sample an amide I peak from the Bcl-2 carbonyls (predominantly) was also observed during SLB fabrication. Once the SLB was formed Bax was injected into the solid/liquid flow cell and its accumulation at the near surface region was monitored through the appearance of a protein Amide I peak.

### Section 2: Additional Information

**Table S1. Scattering length densities of system components used in the models for fitting and understanding of the neutron reflectivity data. Given cardiolipins low volume fraction in the membrane and the mix of tails in the heart CL used, we used the SLD of h-POPC tails only in our analysis. *Value for 100% deuterated tails given, in our models the actual SLD assumes 95% deuteration. Bax and Bcl-2 values were obtained from the Biomolecular Scattering Length Density Calculator(5).**

| Component | Scattering Length Density (in $H_2O$) $\times 10^{-6}$ Å$^{-2}$ | Scattering Length Density (in $D_2O$) $\times 10^{-6}$ Å$^{-2}$ |
|---|---|---|
| POPC lipid head group | 1.81 | 1.81 |
| h-POPC lipid tails | -0.29 | -0.29 |
| d-POPC lipid tails | 6.85 | 6.85 |
| Cardiolipin lipid head | 4.21 | 4.52 |
| Bcl-2 | 1.90 | 3.20 |
| Bax | 1.85 | 3.05 |
| d-Bax | 5.87 | 7.60 |

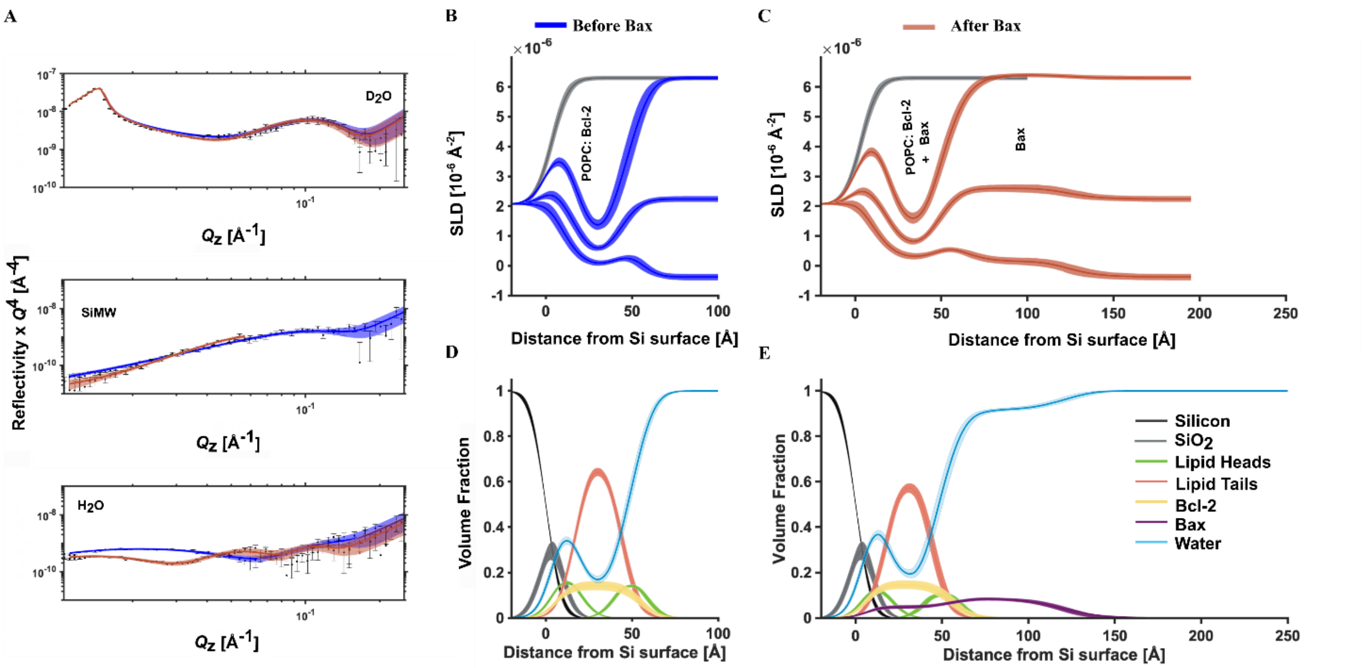


**Figure S1. NR data showing d-Bax binding to a h-POPC Bcl-2 lipid bilayer.** NR data (error bars) and model data fits from a h-POPC: h-Bcl-2 SLB before (blue) and after (red) the interaction of deuterated (d-)Bax are shown in three differing solution isotopic contrast conditions being $D_2O$, Si-MW and $H_2O$ (A) buffer solutions. The scattering length density (SLD) profiles are shown for the surface structure before (B) and after the h-Bax interaction (C). The corresponding component volume fraction profiles are shown before (D) and after (E) the h-Bax interaction as determined from the NR fits. Individual components are color-coded as indicated, with the Bcl-2 distribution in orange and the Bax protein distribution in purple. Line widths in the NR data fits represent the 65% confidence interval of the range of acceptable fits determined from Monte-Carlo-Markov Chain (MCMC) error analysis and the line widths in the SLD and volume fraction profiles represent the ambiguity in the resolved interfacial structure determined from these.

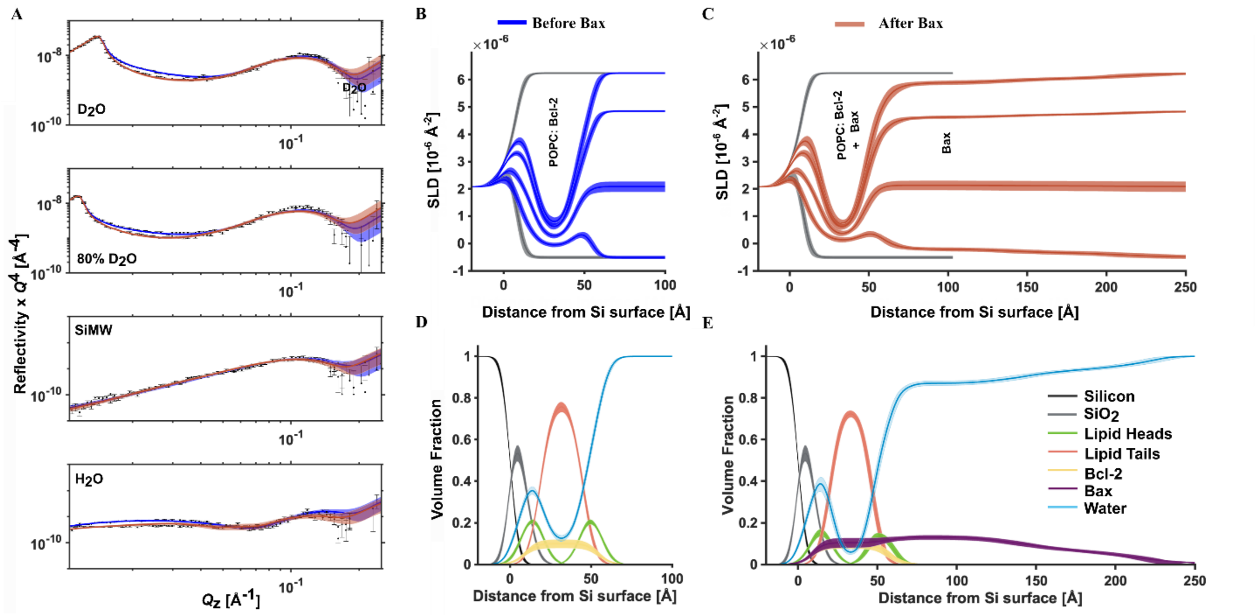


**Figure S2. NR data showing h-Bax binding to a h-POPC Bcl-2 lipid bilayer.** NR data (error bars) and model data fits (lines; s. also supplement Table 1) from a h-POPC/Bcl-2 SLB before (blue) and after (red) the interaction of natural abundance hydrogen (h-)Bax are shown in four different solution isotopic contrast conditions being $D_2O$, 80% $D_2O$, Si-MW and $H_2O$ (A) buffer solutions. The scattering length density (SLD) profiles are shown for the surface structure before (B) and after the h-Bax interaction (C). The corresponding component volume fraction profiles are shown before (D) and after (E) the h-Bax interaction as determined from the NR fits. Individual components are color-coded as indicated, with the Bcl-2 distribution in orange and the Bax protein distribution in purple. Line widths in the NR data fits represent the 65% confidence interval of the range of acceptable fits determined from Monte-Carlo-Markov Chain (MCMC) error analysis and the line widths in the SLD and volume fraction profiles represent the ambiguity in the resolved interfacial structure determined from these.

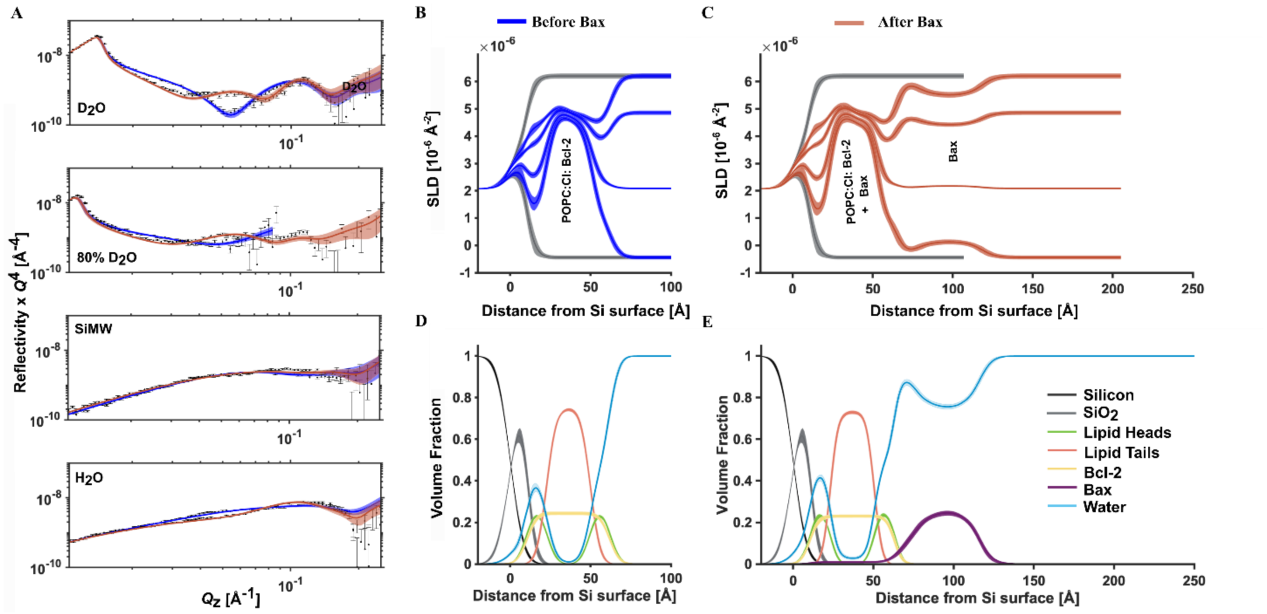


**Figure S3. NR data showing h-Bax binding to a d-POPC cardiolipin (9:1) Bcl2 lipid bilayer.** NR data (error bars) and model data fits from a d-POPC/Bcl-2 SLB before (blue) and after (red) the interaction of natural abundance of hydrogen (h-)Bax are shown in four differing solution isotopic contrast conditions being $D_2O$, Au-MW, Si-MW and $H_2O$ (A) buffer solutions. The scattering length density (SLD) profiles are shown for the surface structure before (B) and after the h-Bax interaction (C). The corresponding component volume fraction profiles are shown before (D) and after (E) the h-Bax interaction as determined from the NR fits. Individual components are color-coded as indicated, with the Bcl-2 distribution in orange and the Bax protein distribution in purple. Line widths in the NR data fits represent the 65% confidence interval of the range of acceptable fits determined from Monte-Carlo-Markov Chain (MCMC) error analysis and the line widths in the SLD and volume fraction profiles represent the ambiguity in the resolved interfacial structure determined from these.

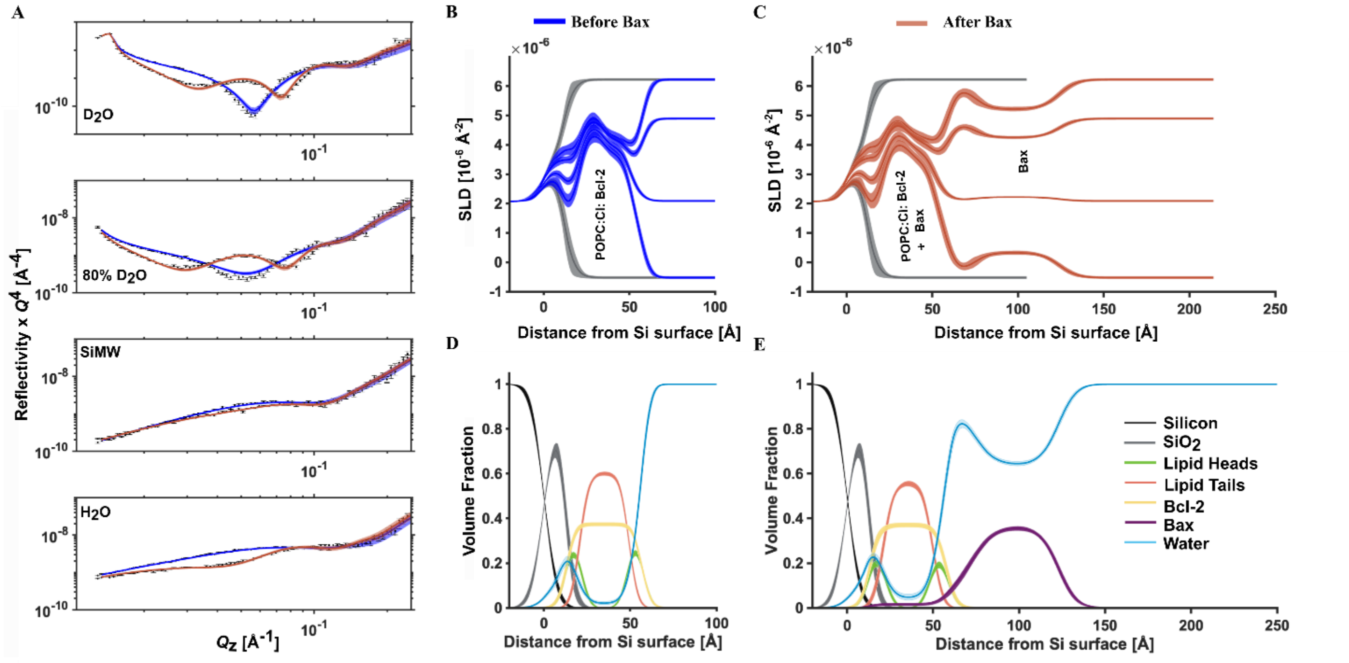


**Figure S4. NR data showing h-Bax binding to a d-POPC h-cardiolipin (9:1) Bcl-2 lipid bilayer.** NR data (error bars) and model data fits from a d-POPC:CL/Bcl-2 SLB before (blue) and after (red) the interaction of natural abundance of hydrogen (h-)Bax are shown in four different solution isotopic contrast conditions being $D_2O$, Au-MW, Si-MW and $H_2O$ (A) buffer solutions. The scattering length density (SLD) profiles are shown for the surface structure before (B) and after the h-Bax interaction (C). The corresponding component volume fraction profiles are shown before (D) and after (E) the h-Bax interaction as determined from the NR fits. Individual components are color-coded as indicated, with the Bcl-2 distribution in orange and the Bax protein distribution in purple. Line widths in the NR data fits represent the 65% confidence interval of the range of acceptable fits determined from Monte-Carlo-Markov Chain (MCMC) error analysis and the line widths in the SLD and volume fraction profiles represent the ambiguity in the resolved interfacial structure determined from these.

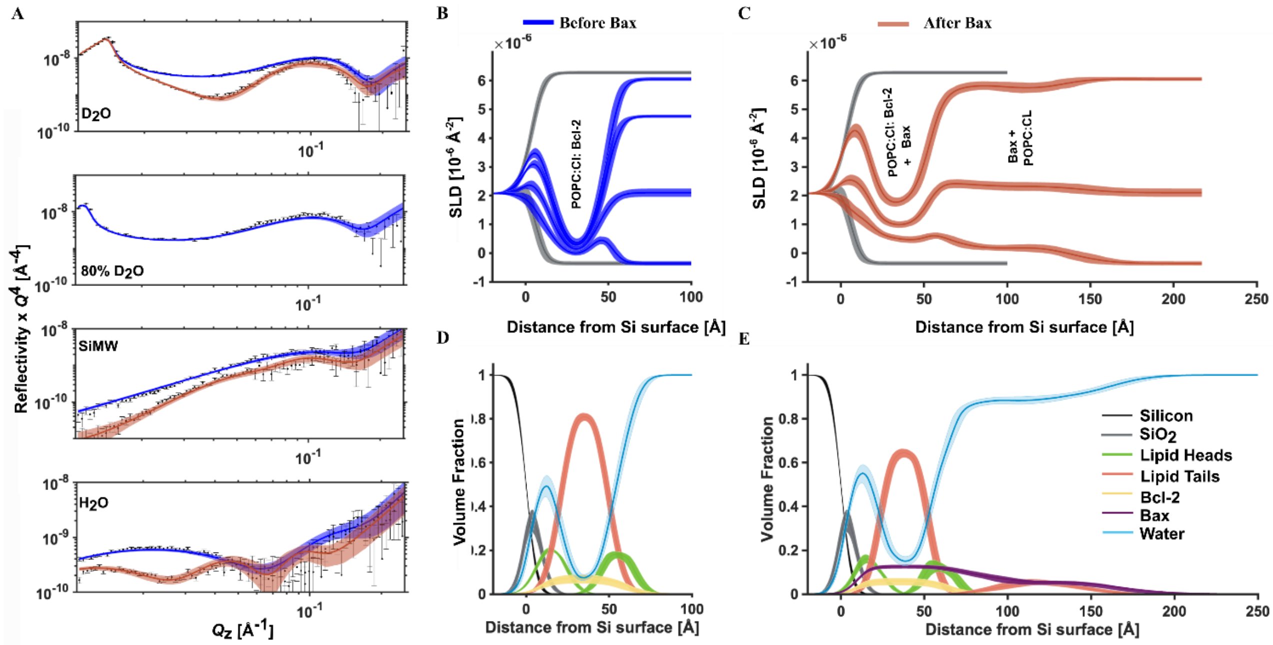


**Figure S5. h-POPC:CL (9:1) Bcl-2 bilayer interaction with h-Bax, at low VF of Bcl-2-lipid removal and poration of the membrane still occurs.** NR data (error bars) and model data fits from a h-POPC:Cl/Bcl-2 SLB before (blue) and after (red) the interaction of natural abundance of hydrogen (h-)Bax are shown in different solution isotopic contrast conditions being $D_2O$, Au-MW, Si-MW and $H_2O$ (A) buffer solutions. The scattering length density (SLD) profiles are shown for the surface structure before (B) and after the h-Bax interaction (C). The corresponding component volume fraction profiles are shown before (D) and after (E) the h-Bax interaction as determined from the NR fits. Individual components are color-coded as indicated, with the Bcl-2 distribution in orange and the Bax protein distribution in purple. Line widths in the NR data fits represent the 65% confidence interval of the range of acceptable fits determined from Monte-Carlo-Markov Chain (MCMC) error analysis and the line widths in the SLD and volume fraction profiles represent the ambiguity in the resolved interfacial structure determined from these.

**Table S2: The resolved structural components before and after the interaction of Bax with SLBs composed of POPC-cardiolipin containing Bcl-2 protein.**

*Values in parentheses represent the 65% confidence intervals determined from MCMC resampling of the experimental data fits.

| | | Lipid Area per Molecule / Å² | Tails Thickness / Å | Tails Composition | Head Group Thickness / Å | Outer Head-group Composition | Membrane Surface Bax/Lipid Complex Thickness / Å | Bax Surface Layer Composition |
|---|---|---|---|---|---|---|---|---|
| **i) h-POPC: CL: Bcl-2** | | 65.7 (64.7 66.8) | 28.4 (27.9 28.8) | Lipid 93.0 (91.0 94.8)<br>Bcl-2 protein 6.9 (5.1 9.0)<br>Solution 0.1 (0.0 0.1) | 6.8 (6.2 7.3) | Lipid 68.5 (62.7 74.6)<br>Bcl-2 protein 6.9 (5.1 9.0)<br>Solution 25.7 (19.2 31.6) | | |
| | **+ d-Bax** | 53.3 (52.2 54.9) | 35.0 (34.0 35.7) | Lipid 68.6 (66.9 70.4)<br>Bcl-2 protein 5.7 (4.1 7.4)<br>Bax protein 12.6 (11.7 13.4)<br>Solution 11.5 (9.5 13.3) | 14.4 (12.4 16.3) | Lipid 36.6 (33.2 40.8)<br>Bcl-2 protein 5.7 (4.1 7.4)<br>Bax protein 12.6 (11.7 13.4)<br>Solution 45.0 (39.2 49.7) | **1,** 34.2 (28.6 40.4)<br>**2,** 42.1 (36.9 46.7)<br>**3,** 34.2 (28.6 40.4)<br>**Total: 109** (100 119) | **1,** Protein 4.6 (3.4 5.8)<br>Solution 95.4 (94.2 96.6)<br>**2,** Protein 8.9 (7.9 10.0)<br>Lipid 24.2 (22.5 25.7)<br>Solution 67.2 (65.1 69.4)<br>3, Protein 4.6 (3.4 5.8)<br>Solution 95.4 (94.2 96.6) |
| **ii) d-POPC: Cl: Bcl-2** | | 68.3 (67.6 69.1) | 27.3 (27.0 27.6) | Lipid 74.8 (74.0 75.5)<br>Bcl-2 protein 24.5 (23.7 25.3)<br>Solution 0.5 (0.4 0.7) | 12.0 (11.7 12.4) | Lipid 30.0 (29.2 30.9)<br>Bcl-2 protein 24.5 (23.7 25.3)<br>Solution 60.4 (59.3 61.7) | | |
| | **+ h-Bax** | 64.7 (63.9 65.6) | 28.8 (28.4 29.2) | Lipid 73.1 (72.2 74.0)<br>Bcl2 protein 24.5 (23.7 25.3)<br>Bax protein 0.7 (0.14 1.68)<br>Solution 2.6 (2.0 3.3) | 12.7 (12.3 13.0) | Lipid 27.9 (26.0 29.9)<br>Bcl-2 protein 24.5 (9.6 13.1)<br>Bax protein 0.7 (4.1 5.6)<br>Solution 46.3 (45.2 47.5) | **1,** 12.3 (10.7 14.2)<br>**2,** 40.1 (37.4 42.9)<br>**Total: 53** (51 54) | **1,** Protein 9.0 (8.0 10.0)<br>Solution 91.0 (90.0 92.0)<br>**2,** Protein 25.0 (23.6 26.4)<br>Solution 75.0 (73.6 76.4) |
| **iii) d-POPC: Cl: Bcl-2** | | 65.9 (64.5 67.2) | 28.3 (27.8 28.9) | Lipid 59.3 (58.2 60.3)<br>Bcl-2 protein 38.2 (37.4 39.0)<br>Solution 2.1 (1.4 2.69) | 7.4 (7.1 7.8) | Lipid 40.2 (38.2 42.3)<br>Bcl-2 protein 38.2 (37.4 39.0)<br>Solution 34.0 (30.5 37.9) | - | - |
| | **+ h-Bax** | 65.1 (63.3 67.1) | 28.6 (27.7 29.5) | Lipid 56.2 (54.9 57.5)<br>Bcl-2 protein 37.1 (36.0 38.1)<br>Bax protein 1.3 (0.8 2.0)<br>Solution 4.4 (3.1 5.9) | 7.9 (7.1 8.4) | Lipid 56.2 (54.9 57.5)<br>Bcl-2 protein 37.1 (36.0 38.1)<br>Bax protein 1.3 (0.8 2.0)<br>Solution 24.5 (21.9 27.0) | **1,** 15.5 (13.9 17.1)<br>**2,** 49.5 (47.2 52.1)<br>**Total: 65** (64 67) | **1,** Protein 6.3 (4.2 8.5)<br>Solution 93.7 (91.5 96.7)<br>**2,** Protein 35.8 (34.7 36.8)<br>Solution 64.2 (63.2 65.3) |

**Table S3. The resolved structural components before and after the interaction of Bax with SLBs composed of POPC containing Bcl-2 protein.**
*Values in parentheses represent the 65% confidence intervals determined from MCMC resampling of the experimental data fits.

| | | **Average Lipid Area per Molecule / Å²** | **Tails Thickness / Å** | **Tails Composition** % | **Head Group Thickness/ Å** | **Outer Head-group Composition** % | **Membrane Surface Bax/Lipid Complex Thickness / Å** | **Bax Surface Layer Composition** % |
|---|---|---|---|---|---|---|---|---|
| **I)**<br>**h-POPC: Bcl-2** | | 72.4 (70.7 74.3) | 25.7 (25.1 26.4) | Lipid 79.8 (76.8 82.8)<br>Bcl-2 protein 9.9 (9.5 10.1)<br>Solution 10.2 (8.5 11.9) | 10.2 (9.8 10.6) | Lipid 35.5 (34.2 37.2)<br>Bcl-2 protein 9.9 (9.5 10.1)<br>Solution 64.8 (63.2 66.5) | | |
| | **+ h-Bax** | 67.8 (66.1 69.7) | 27.5 (26.8 28.2) | Lipid 76.7 (74.7 78.8)<br>Bcl-2 protein 8.3 (6.3 10.1)<br>Bax protein 10.4 (8.0 12.7)<br>Solution 2.3 (1.6 3.2) | 12.9 (11.6 13.9) | Lipid 38.8 (35.9 42.2)<br>Bcl-2 protein 8.3 (6.3 10.1)<br>Bax protein 10.4 (8.0 12.7)<br>Solution 50.3 (48.2 52.6) | **1,** 88.4 (78.0 98.9)<br>**2,** 80.8 (72.6 87.6)<br>**Total:** 163 (154 174) | **1,** Protein 13.0 (11.9 14.0)<br>Solution 87.0 (85.9 88.2)<br>**2,** Protein 5.8 (4.9 6.7)<br>Solution 94.2 (93.3 95.1) |
| **Ii)**<br>**h-POPC: Bcl-2** | | 73.0 (70.6 75.4) | 25.5 (24.7 26.4) | Lipid 76.3 (73.6 79.2)<br>Bcl-2 protein 14.2 (12.3 16.2)<br>Solution 11.2 (9.4 11.9) | 11.8 (11.4 12.3) | Lipid 29.3 (28.2 30.5)<br>Bcl-2 protein 14.2 (12.3 16.2)<br>Solution 56.2 (52.1 59.5) | | |
| | **+ d-Bax** | 63.7 (61.6 65.8) | 29.3 (28.3 30.3) | Lipid 69.4 (67.1 71.9)<br>Bcl-2 protein 11.3 (9.6 13.1)<br>Bax protein 4.8 (4.1 5.6)<br>Solution 11.3 (9.6 13.0) | 14.3 (13.5 15.1) | Lipid 37.8 (36.3 39.6)<br>Bcl-2 protein 11.3 (9.6 13.1)<br>Bax protein 4.8 (4.1 5.6)<br>Solution 45.7 (42.1 47.4) | **1,** 31.6 (23.4 40.6)<br>**2,** 37.9 (30.5 45.0)<br>**Total:** 68 (63 74) | **1,** Protein 9.0 (8.0 10.0)<br>Solution 91.0 (90.0 92.0)<br>**2,** Protein 4.5 (3.5 5.8)<br>Solution 95.5 (94.2 96.5) |
| **Iii)**<br>**d-POPC: Bcl-2** | | 76.5 (73.2 79.9) | 24.4 (23.3 25.5) | Lipid 48.6 (45.5 51.7)<br>Bcl-2 protein 39.9 (38.5 41.3)<br>Solution 11.5 (9.4 13.6) | 9.3 (8.6 10.0) | Lipid 22.5 (21.2 24.0)<br>Bcl-2 protein 39.9 (38.5 41.3)<br>Solution 65.0 (60.9 67.8) | - | - |
| | **+ h-Bax** | 81.3 (77.9 84.7) | 22.9 (22.0 23.9) | Lipid 46.0 (42.7 49.1)<br>Bcl-2 protein 37.3 (35.6 39.3)<br>Bax protein 9.4 (7.1 11.7)<br>Solution 4.5 (3.0 6.1) | 8.8 (8.1 9.6) | Lipid 39.0 (35.9 42.6)<br>Bcl-2 protein 37.3 (35.6 39.3)<br>Bax protein 9.4 (7.1 11.7)<br>Solution 50.7 (48.0 53.0) | **1,** 55 (53.4 56.7)<br>**2,** 52.7 (41.1 63.4)<br>**Total:** 108 (95 120) | **1,** Protein 26.7 (24.9 27.4)<br>Solution 73.8 (72.6 75.1)<br>**2,** Protein 4.5 (3.5 5.8)<br>Solution 95.5 (94.2 96.5) |

**Table S4. The time-resolved structural components during the interaction of Bax with SLB composed of d-POPC containing Bcl-2 protein. Structural components on the data set before and after Bax interaction shown in Table S3 part iii). Data sets measured in D2O subphase contrast only and parameters fixed other than the volume fraction and thickness parameters for Bax.**
*Values in parentheses represent the 65% confidence intervals determined from MCMC resampling of the experimental data fits, note that when a prior has a hard limit, such as the relative multipliers below having a limit of 1, then the distribution of the posterior used for confidence interval estimation can be non-gaussian and the best fit value can lie slightly outside the 65% C.I.

| Time since Bax injection | K | K_d | VF bilayer/% | VF first layer /% | VF second layer /% | Thickness of Bax layer 1/Å | Thickness of Bax layer 2/Å |
|---|---|---|---|---|---|---|---|
| 0 | 0 | 0 | 0 | 0 | 0 | 0 | 0 |
| 47-83 | 0.47 (0.37 0.80) | 0.43 (0.29 0.71) | 4.7 (3.7 8.0) | 12.3 (9.6 20.9) | 2.1 (1.7 3.6) | 23.7 (20.2 39.1) | 22.7 (15.3 37.4) |
| 83-120 | 0.59 (0.45 0.84) | 0.49 (0.36 0.75) | 5.9 (4.4 8.4) | 15.5 (11.7 22.1) | 2.7 (2.0 3.8) | 27.0 (24.6 41.2) | 25.9 (18.8 39.4) |
| 120-156 | 0.63 (0.48 0.84) | 0.59 (0.44 0.80) | 6.3 (4.8 8.3) | 16.5(12.7 21.9) | 2.8 (2.2 3.8) | 32.5 (26.7 44.1) | 31.2 (23.1 42.3) |
| 156-193 | 0.55 (0.44 0.75) | 0.99 (0.50 0.92) | 5.4 (4.4 7.4) | 14.3(11.6 19.6) | 2.5 (2.0 3.4) | 54.5 (24.4 50.4) | 52.3 (26.4 48.3) |
| 193-234 | 0.67 (0.56 0.84) | 0.81 (0.59 0.90) | 6.6 (5.5 8.4) | 17.4(14.6 22.1) | 3.0 (2.5 3.8) | 44.4 (30.6 49.8) | 42.5 (30.9 47.7) |
| 234-271 | 0.70 (0.58 0.85) | 0.85 (0.60 0.92) | 6.9 (5.8 8.5) | 18.2(15.2 22.3) | 3.1(2.6 3.8) | 47.1 (32.0 50.5) | 45.1 (31.7 48.4) |
| 271-307 | 0.78 (0.64 0.91) | 0.80 (0.64 0.91) | 7.7 (6.4 9.1) | 20.3(16.7 23.8) | 3.5(2.9 4.1) | 43.8 (35.2 50.2) | 41.9 (33.6 48.2) |
| 307-343 | 0.79 (0.63 0.90) | 0.85 (0.64 0.92) | 7.8 (6.3 8.9) | 20.6(16.6 23.4) | 3.5(2.8 4.0) | 46.6 (34.9 50.7) | 44.6 (34.0 48.6) |
| 343-380 | 0.82 (0.69 0.92) | 0.88 (0.71 0.94) | 8.1(6.8 8.9) | 21.4(18.0 24.1) | 3.7(3.1 4.1) | 48.2 (37.8 51.7) | 46.2 (37.6 49.5) |
| 380-416 | 0.86(0.72 0.93) | 0.87(0.70 0.93) | 8.5(7.2 9.2) | 22.4(18.8 24.4) | 3.8(3.2 4.2) | 47.7 (39.6 51.4) | 45.7 (36.9 49.2) |
| 416-453 | 0.84 (0.70 0.92) | 0.94 (0.74 0.95) | 8.3(6.9 9.3) | 21.9(18.2 24.2) | 3.8(3.1 4.1) | 51.5 (38.3 52.3) | 49.4 (39.0 50.2) |
| 453-489 | 0.89 (0.74 0.94) | 1.00 (0.78 0.96) | 8.8(7.4 9.2) | 23.2(19.4 24.7) | 4.0(3.3 4.2) | 54.8 (40.8 52.9) | 52.5 (41.3 50.7) |
| 489-526 | 0.93 (0.77 0.95) | 1.00(0.79 0.96) | 9.2(7.7 9.4) | 24.2(20.1 25.0) | 4.2(3.5 4.3) | 54.8 (42.4 53.1) | 52.5 (41.5 50.8) |

| | | | | | | | |
|---|---|---|---|---|---|---|---|
| 526-563 | 0.92 (0.78 0.96) | 1.00 (0.82 0.97) | 9.1(7.8 9.5) | 24.0(20.4 25.0) | 4.1(3.5 4.3) | 55.0 (43.0 53.4) | 52.7 (43.2 51.2) |
| 563-603 | 0.96 (0.80 0.96) | 0.97 (0.80 0.96) | 9.5(7.9 9.5) | 25.1(20.9 25.2) | 4.3(3.6 4.3) | 53.2 (43.9 53.4) | 51.0 (42.4 50.9) |
| 603-641 | 1.00 (0.80 0.96) | 1.00 (0.82 0.97) | 9.9(7.9 9.6) | 26.1(20.9 25.2) | 4.5(3.6 4.3) | 55.0 (44.0 53.1) | 52.7 (43.2 51.2) |
| 641-693 | 0.95 (0.83 0.97) | 1.00 (0.86 0.98) | 9.4(8.3 9.6) | 24.8(21.9 25.4) | 4.3(3.8 4.4) | 55.0 (46.0 53.4) | 52.7 (45.6 51.6) |
| 693-738 | 0.97 (0.84 0.97) | 1.00 (0.86 0.98) | 9.6(8.3 9.7) | 25.3(22.0 25.4) | 4.4(3.8 4.4) | 55.0 (46.2 53.9) | 52.7 (45.2 51.6) |
| Final structure | 1 | 1 | 10.0% | 26.2 | 4.5% | 55.1 Å | 52.8 Å |

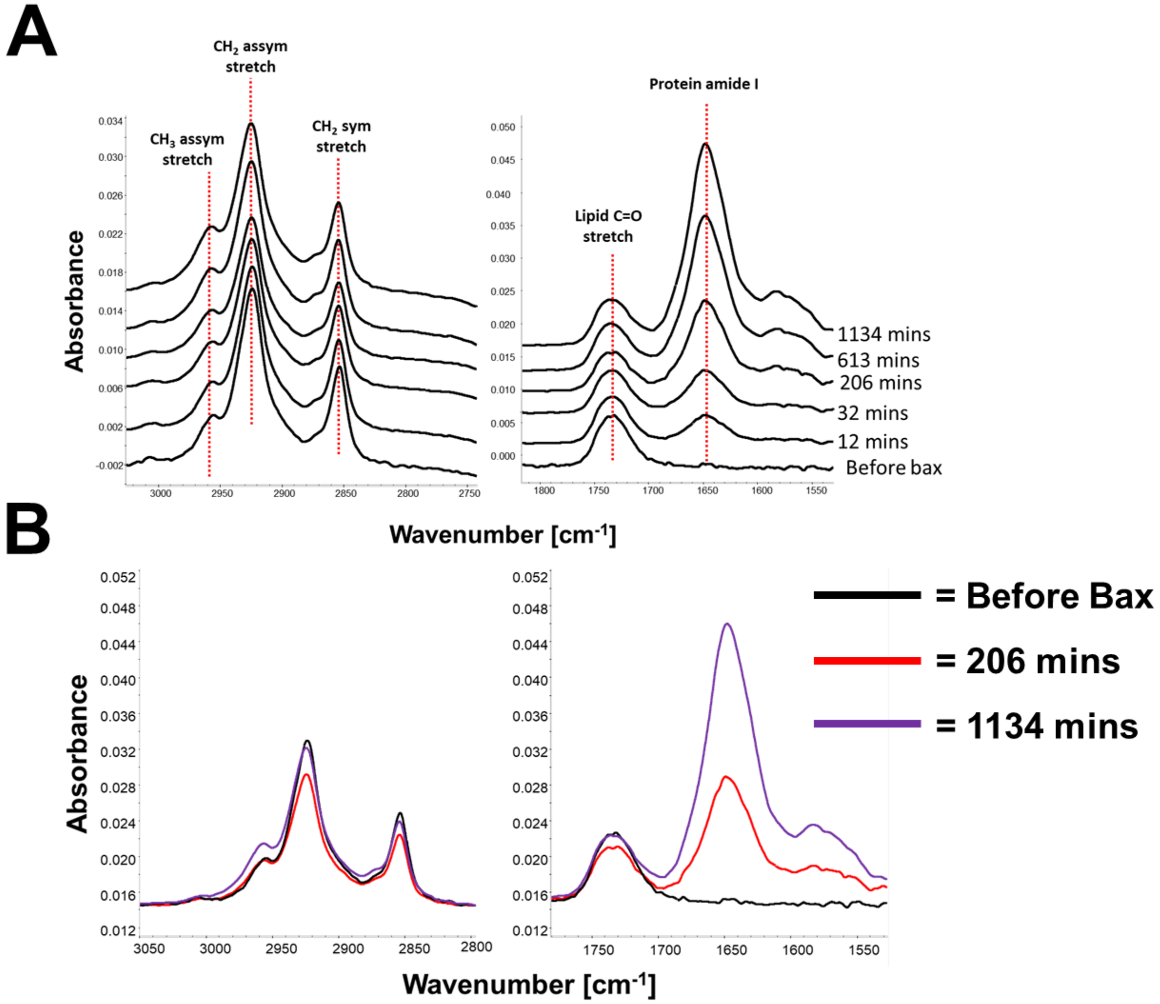


**Figure S6. ATR-FTIR data obtained during the interaction of h-Bax with a h-POPC SLB.** Changes in the CH stretch (A, Left) and Amide I (A, right) regions of the spectra are shown against time. An overlaid comparison of these regions is given; (B) showing the accumulation of Bax at the SLB coated surface and concurrent changes in the CH2 stretches.

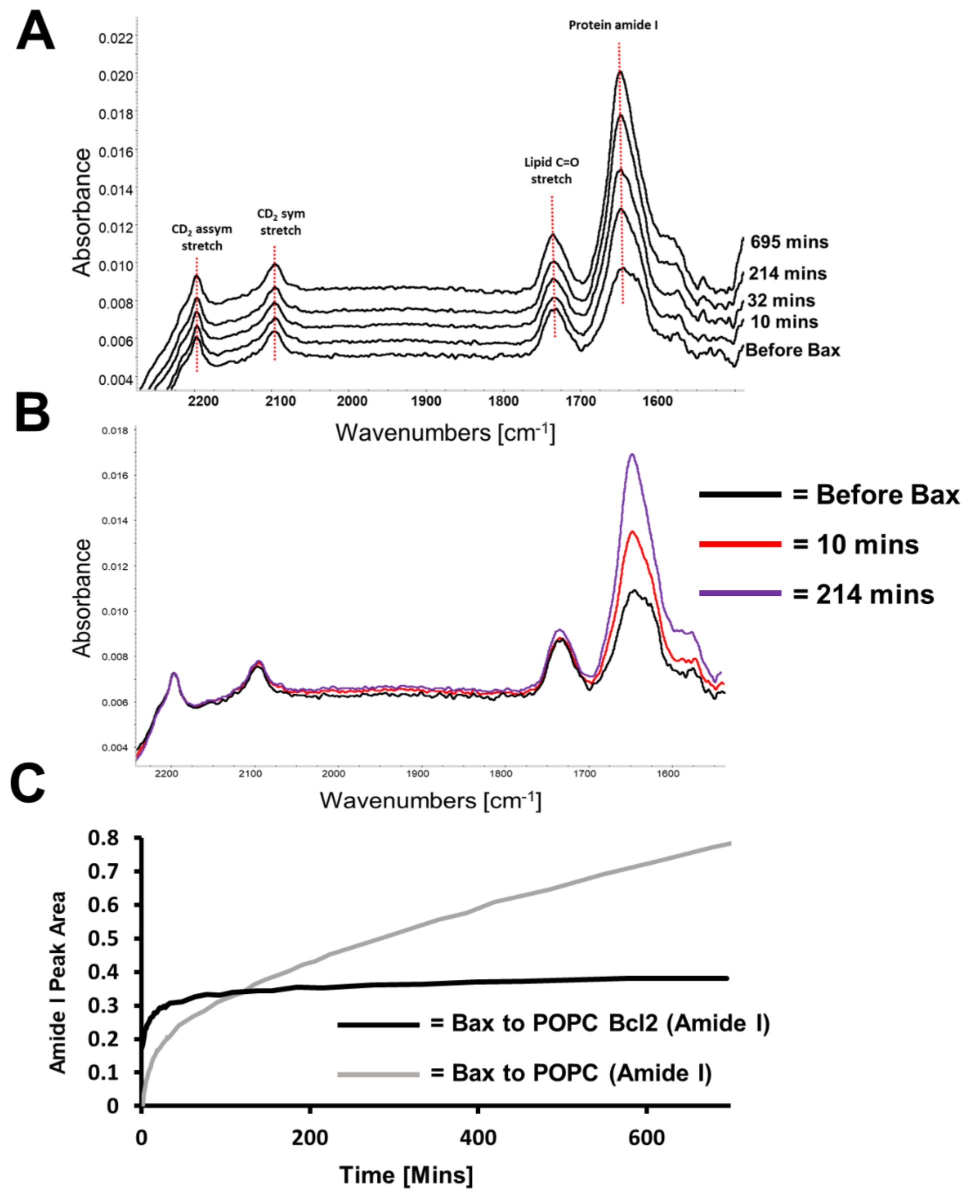


**Figure S7. ATR-FTIR data from the interaction of h-Bax with a Bcl-2/d-POPC SLB.** Changes in the CD stretching (lipid tail CD2 and CD3) and Amide I (Bcl-2 and Bax protein) region of the spectra against time are shown (A). An overlaid comparison of the spectra at differing times during the Bax binding process showing an increase in the amide I peak intensity no significant changes in the CD stretching region concurrent with Bax binding to, rather than disruption of the Bcl-2/POPC SLB (B). A comparison between the increase in the Amide I (protein carbonyl) band due to Bax binding to the POPC online (C, back line) and Bcl-2/POPC (C, grey line) SLBs, indicating that Bax binding to a Bcl-2 containing surface occurs on a faster time scale than Bax induced pore formation.

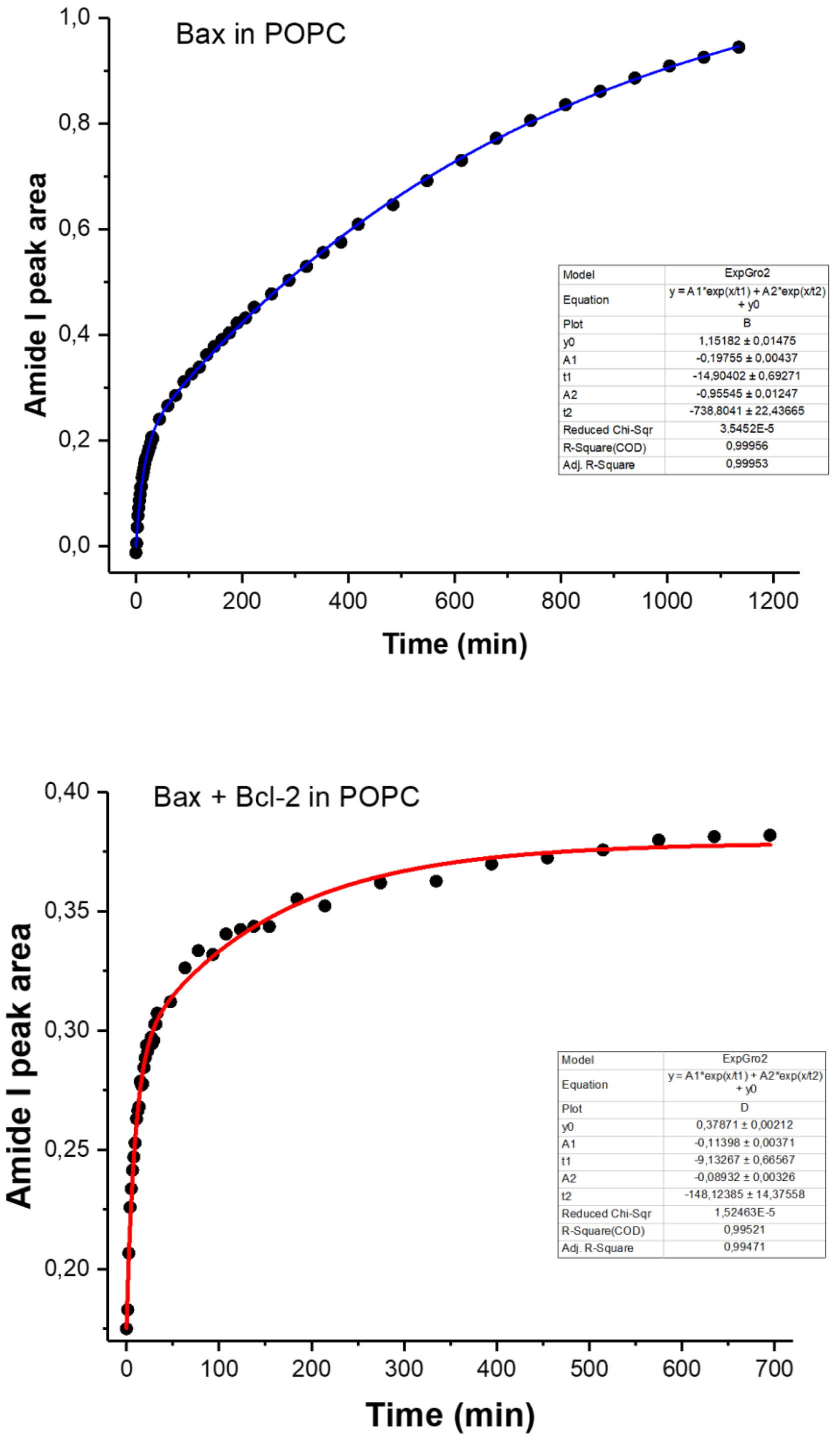


**Figure S8.** ATR-FTIR data showing the increase in Amide 1 peak area after addition of Bax to a SSB of POPC (top) and a SSB of POPC containing Bcl-2 (bottom) respectively. Collection times for each ATR-FTIR dataset were ~80s. Fits to exponential models show that association of Bax to these bilayers is a two-step process. The Bax association time constants for POPC bilayers were t1=-14.9 min and t2=738.8 min, and for POPC-Bcl-2 bilayer t1=-9.1 min and t2=148.1 min.

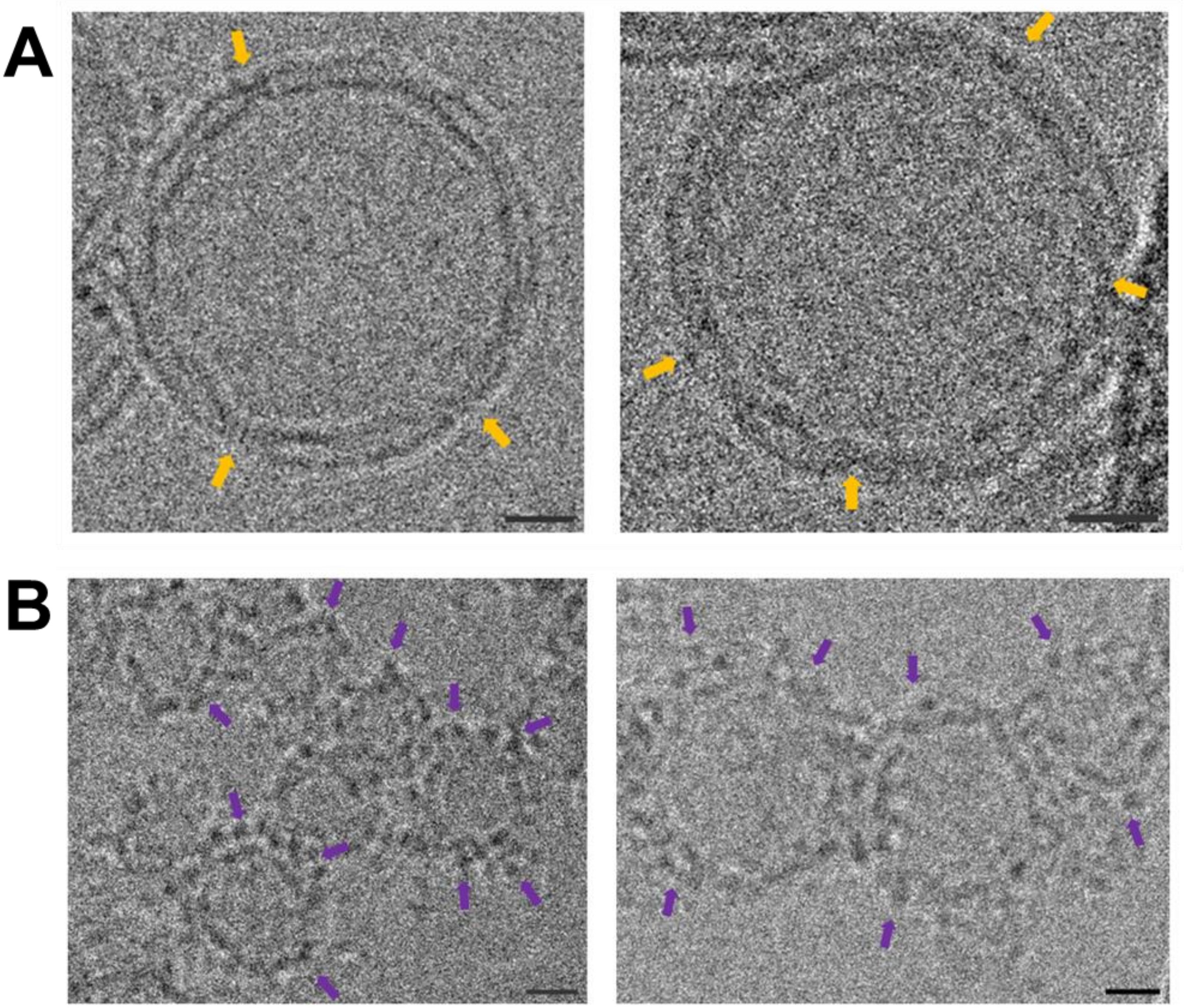


**Figure S9. Selected EM microscopy image regions showing the presence of Bcl-2 (A, yellow arrows) and Bax (B, purple arrows) within and on the surface of h-Bcl-2 : d-POPC vesicles respectively.** Images show d-POPC : h-Bcl-2 without (A) and after (B) the incubation in the presence of h-Bax for 60 minutes. The scale bar at the bottom right of each panel is 10 nm.

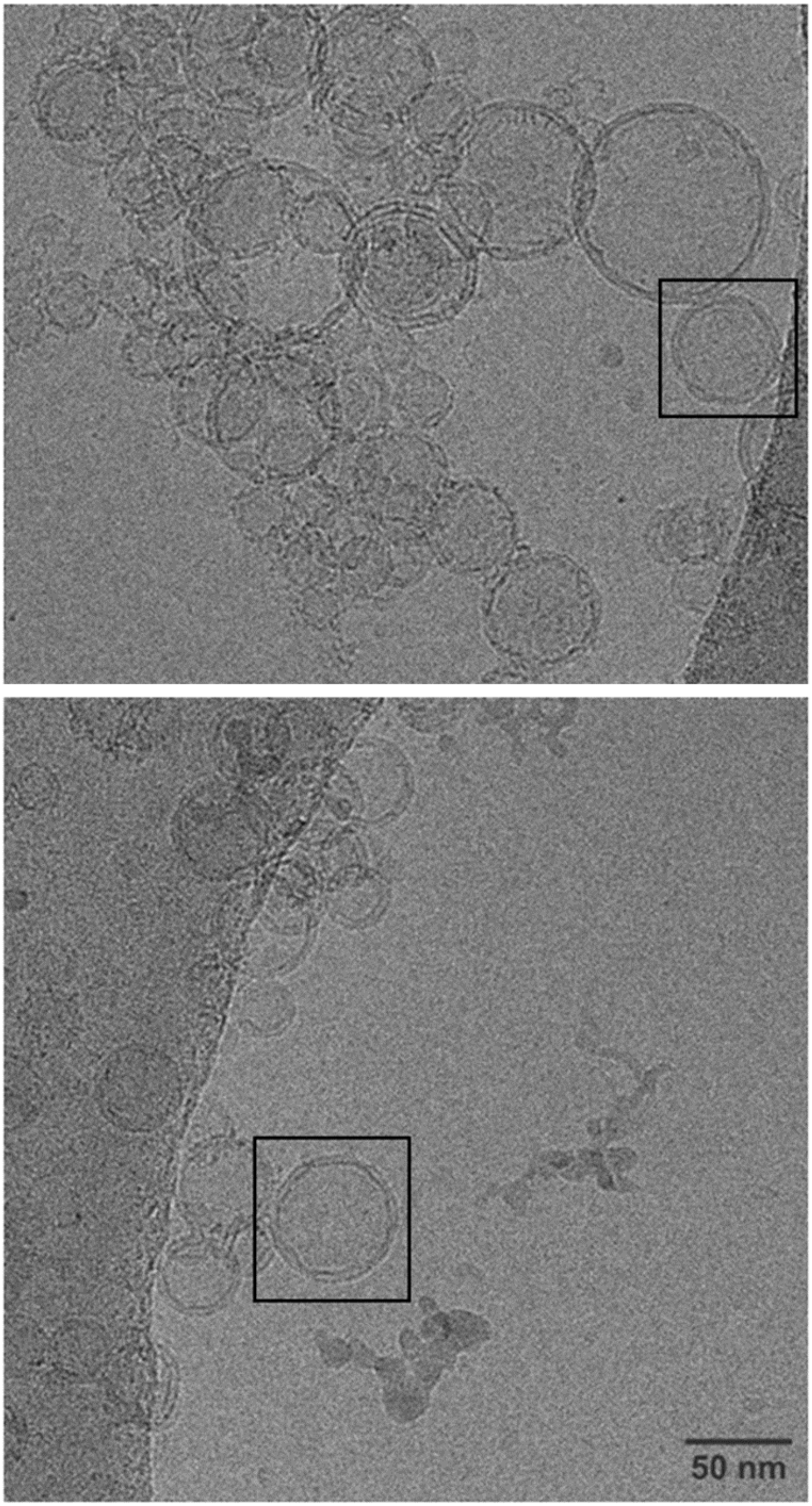


**Figure S10.** EM micrographs of d-POPC : h-Bcl-2 vesicles. Areas shown in black squares are shown expanded in Fig 2 and SI Fig 10.

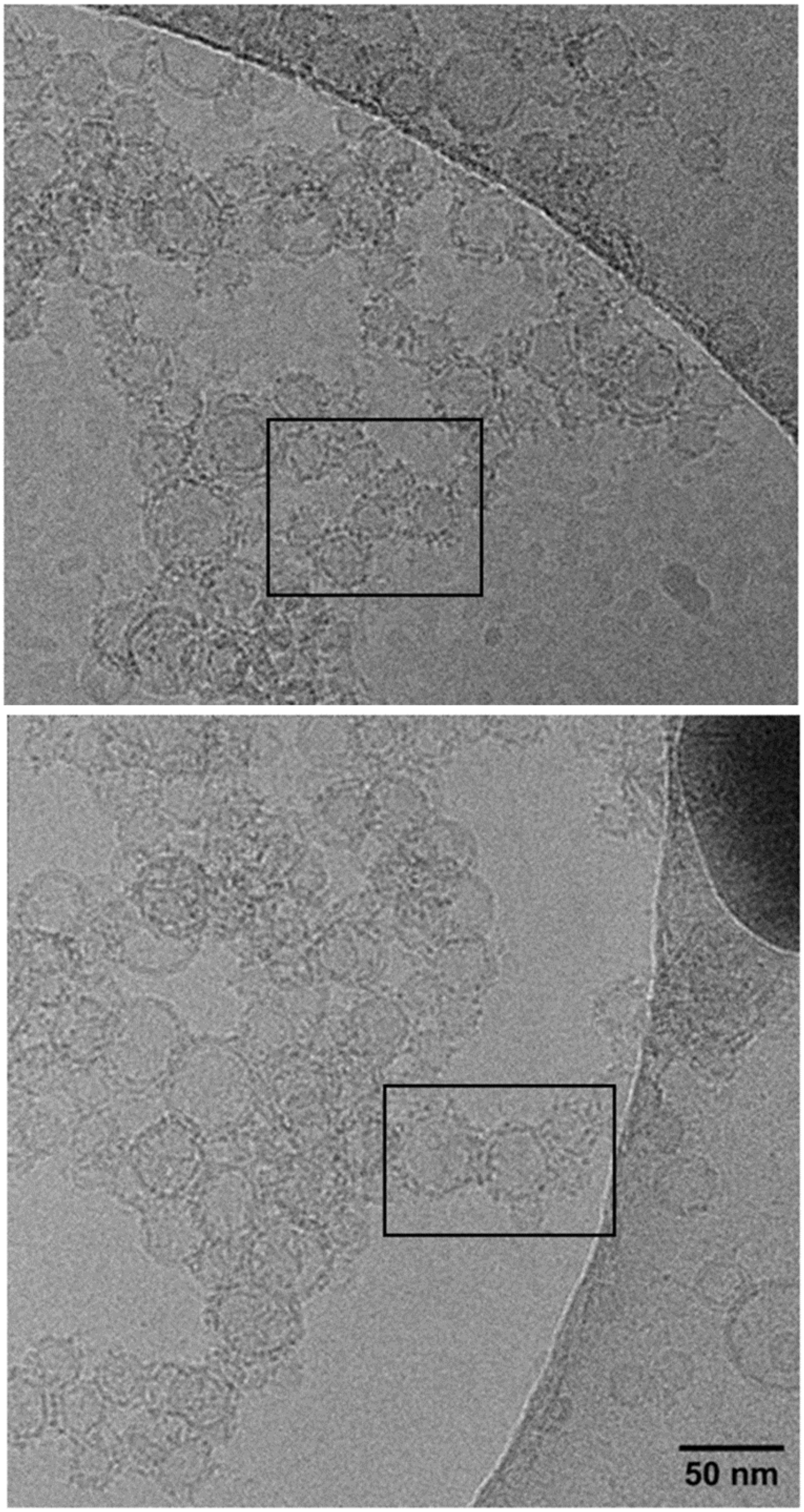


**Figure S11.** EM micrographs of h-Bax bound d-POPC : h-Bcl-2 vesicles. Areas shown in black squares are shown expanded in Fig 2 and SI Fig 10.

**Section 3: Additional Modelling**

**Modelling of Bax dimer structure for comparison with NR resolved Membrane Surface Bax distributions**

Potential Bax dimer configurations were generated using AlphaFold2-multimer (*6*), making use of ColabFold v1.5.5 (*7*). The input protein sequence for Bax was obtained from UniProtKB assession number Q07812. Alphafold was set to detect potential templates (*8*) in the pdb100 database (*9*)) and perform AMBER relaxation (*10*) on the top 5 ranked structures. The radius (and subsequent diameter) of gyration was evaluated for each of the 5 relaxed structures. Sequence coverage was 100% for all positions. The predicted local distance difference test (pLDDT) scores and predicted aligned error (PAE) are shown in figure S13. As might be expected, the N-terminal residues show some of the highest uncertainty in position.

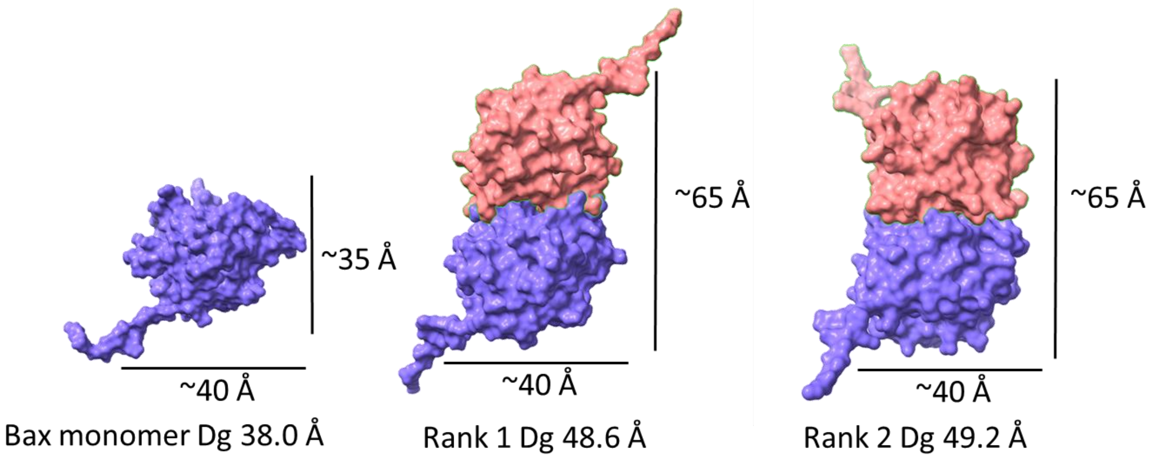


**Figure S12. Bax monomer structure (PDB 1F16) and top two ranked dimer models with relative structural length scales**.

The radii of gyration obtained from these top ranked models are highly comparable to the distances of the Bax distributions indicated from the reflectometry analysis where the individual Bax distributions were found to be 52 – 68 Å in length in the cases where discernible protein distributions were observed on the membrane surface (see main manuscript Table 2). Figure S12 shows a comparison of a Bax monomer with the top 2 ranked dimer structures from AlphaFold-multimer modelling. In all cases the dimers showed a prolate structure with a minor axis of ~40 Å and a major axis of ~65 Å (not including the N-terminal tail regions). As the non-disruptive Bax distributions found on the membrane surfaces were

52 – 68 Å in length this suggests the Bcl-2 bound Bax distributions measured in the vertical direction across the membrane surface were upright or tilted Bax dimers configurations.

**Table S5. Radius of gyration of top 5 relaxed structures from AlphaFold.**

| Structure Rank | Radius of gyration (Å) | Diameter of gyration (Å) |
|---|---|---|
| Monomer structure pdb 1F16 | 19.0 | 38.0 |
| Dimer Rank 1 | 24.3 | 48.6 |
| Dimer Rank 2 | 24.6 | 49.2 |
| Dimer Rank 3 | 24.9 | 49.8 |
| Dimer Rank 4 | 24.5 | 49.0 |
| Dimer Rank 5 | 24.7 | 49.4 |

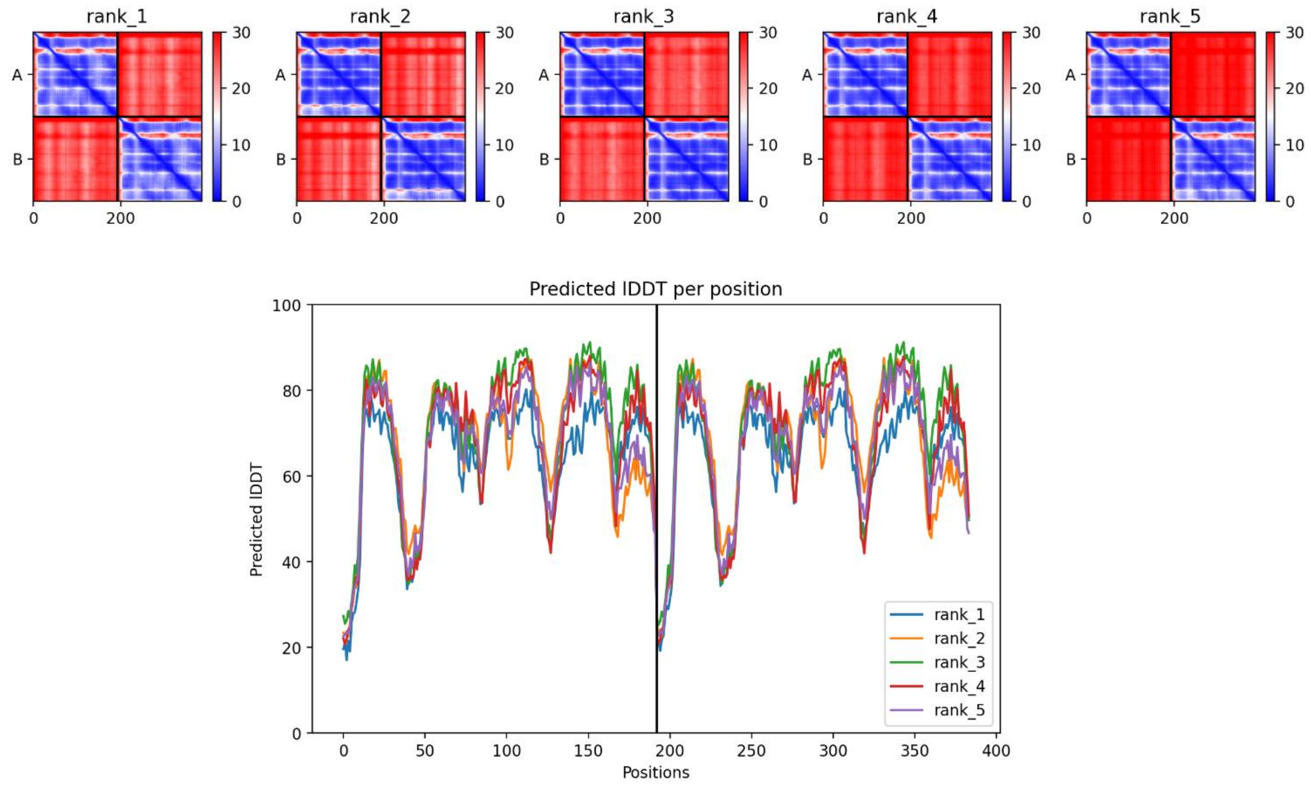


**Figure S13. Predicted aligned error and pLDDT scores for AlphaFold prediction**
